# The Evolution of White Matter Changes After Mild Traumatic Brain Injury: A DTI and NODDI Study

**DOI:** 10.1101/345629

**Authors:** Eva M. Palacios, Julia P Owen, Esther L. Yuh, Maxwell B. Wang, Mary J. Vassar, Adam R. Ferguson, Ramon Diaz-Arrastia, Joseph T. Giacino, David O. Okonkwo, Claudia S. Robertson, Murray B. Stein, Nancy Temkin, Sonia Jain, Michael McCrea, Christine L. Mac Donald, Harvey S. Levin, Geoffrey T. Manley, Pratik Mukherjee, the TRACK-TBI Investigators

## Abstract

Neuroimaging biomarkers show promise for improving precision diagnosis and prognosis after mild traumatic brain injury (mTBI), but none has yet been adopted in routine clinical practice. Biophysical modeling of multishell diffusion MRI, using the neurite orientation dispersion and density imaging (NODDI) framework, may improve upon conventional diffusion tensor imaging (DTI) in revealing subtle patterns of underlying white matter microstructural pathology, such as diffuse axonal injury (DAI) and neuroinflammation, that are important for detecting mTBI and determining patient outcome. With a cross-sectional and longitudinal design, we assessed structural MRI, DTI and NODDI in 40 mTBI patients at 2 weeks and 6 months after injury and 14 matched control participants with orthopedic trauma but not suffering from mTBI at 2 weeks. Self-reported and performance-based cognitive measures assessing postconcussive symptoms, memory, executive functions and processing speed were investigated in post-acute and chronic phase after injury for the mTBI subjects. Machine learning analysis was used to identify mTBI patients with the best neuropsychological improvement over time and relate this outcome to DTI and NODDI biomarkers. In the cross-sectional comparison with the trauma control group at 2 weeks post-injury, mTBI patients showed decreased fractional anisotropy (FA) and increased mean diffusivity (MD) on DTI mainly in anterior tracts that corresponded to white matter regions of elevated free water fraction (FISO) on NODDI, signifying vasogenic edema. Patients showed decreases from 2 weeks to 6 months in white matter neurite density on NODDI, predominantly in posterior tracts. No significant longitudinal changes in DTI metrics were observed. The machine learning analysis divided the mTBI patients into two groups based on their recovery. Voxel-wise group comparison revealed associations between white matter orientation dispersion index (ODI) and FISO with degree and trajectory of improvement within the mTBI group. In conclusion, white matter FA and MD alterations early after mTBI might reflect vasogenic edema, as shown by elevated free water on NODDI. Longer-term declines in neurite density on NODDI suggest progressive axonal degeneration due to DAI, especially in tracts known to be integral to the structural connectome. Overall, these results show that the NODDI parameters appear to be more sensitive to longitudinal changes than DTI metrics. Thus, NODDI merits further study in larger cohorts for mTBI diagnosis, prognosis and treatment monitoring.

## INTRODUCTION

Despite increasing evidence from preclinical and human studies that mild traumatic brain injury (mTBI) causes axonal shearing injury of white matter microstructure that can affect the longterm cognitive, neuropsychiatric, and social domains of function, the lack of reliable objective tools to measure such pathology is a barrier to clinical translation (Manley & Mass, 2013 JAMA; Bigler a al.2013, Levin & Diaz-Arrastia, 2015 Lancet; Radhakrishnan et al., 2016 Lancet). The common assumption, even among healthcare professionals, those mTBI patients will return to premorbid levels of function shortly after the traumatic event often results in these patients not receiving appropriate follow-up care after the acute injury. Sensitive tests that can detect white matter microstructural alterations early after injury and inform the likely path to recovery are needed to improve care and to develop future therapies.

Diffusion tensor imaging (DTI) is the most extensively used technique worldwide to study the microstructural properties of white matter in vivo (Basser et al., 1994; Mori et al., 2006; Mukherjee et al., 2008). DTI studies of mTBI have shown microstructural white matter disruption that can lead to neurocognitive and behavioral deficits after mTBI (Yuh EL et al., 2014 JN, Croall ID et al., 2014 Neurology; Oehr, et al., 2017 Meta-analysis). However, traditional DTI metrics such as mean diffusivity (MD) and fractional anisotropy (FA) represent basic statistical descriptions of diffusion that do not directly correspond to biophysically meaningful parameters of the underlying tissue. Furthermore, DTI assumes Gaussian diffusion within a single microstructural compartment and is therefore insensitive to the complexity of white matter microstructure, which requires a non-Gaussian model with multiple compartments (Jones et al, 2010). Perhaps as a result, prior DTI studies have produced conflicting results with some papers reporting abnormally reduced white matter FA in mTBI and others reporting elevations or no change in FA (Eirud et al., 2014). Other contributing factors to this discordance in the literature include small effect sizes of DTI changes due to mTBI, small sample sizes and the dynamic nature of microstructural white matter alterations after mTBI.

In this investigation, we overcome the limitations of DTI by applying a more advanced multi-compartment diffusion model known as neurite orientation dispersion and density imaging (NODDI) (Zhang et al., 2012; Ojelescu et al., 2017 for review). NODDI leverages recent progress in high-performance magnetic field gradients for MRI scanners that can achieve diffusion-weighting factors much higher than the standard b=1000 s/mm^2^ for DTI and therefore probe more complex non-Gaussian properties of white matter diffusion. The NODDI biophysical model uses this richer diffusion imaging data to measure properties of three microstructural environments: intracellular, extracellular, and free water. One such metric is the intracellular volume fraction, referred to as the neurite density index (NDI) and which primarily represents axonal density within white matter. Another is orientation dispersion index (ODI) of neurites, which is higher in loosely organized white matter and lower in tracts with largely parallel fiber bundles such as the corpus callosum. The volume fraction of the isotropic diffusion compartment (FISO) estimates the free water content (Zhang et al.,2012). These NODDI parameters have been validated in histopathological studies of animal and human brains (Sepehrband, et al., 2015; Sato et al., 2016), been used to study brain development (Yi Shi Chang et al., 2015; Mah et al., 2017) and to detect subtle brain damage in other disorders (Billiet et al., 2015; Timmers et al., 2015; Kamagata et al., 2016; Caverzasi E, et al., 2016; Schneider T et al., 2017).

In this study, we employ NODDI to: i) investigate early white matter changes at two weeks post-injury in mTBI patients versus trauma controls; ii) determine the evolution of white matter changes of the mTBI patients from two weeks to six months after injury; and iii) explore the prognostic significance of these white matter microstructural changes for symptomatic and cognitive outcome after mTBI. Comparing DTI to NODDI serially after mTBI, we hypothesize that the early microstructural white matter changes of mTBI are driven by increases in free water, such as from vasogenic edema due to neuroinflammation, whereas longer-term changes reflect decreases in axonal density due to evolving white matter degeneration.

## MATERIALS AND METHODS

### Participants

All participants were enrolled at the Zuckerberg San Francisco General Hospital and Trauma Center as part of the prospective Transforming Research and Clinical Knowledge in Traumatic Brain Injury project (TRACK-TBI) (Yue et al., 2013). A total sample of 40 mTBI patients (age: 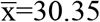 years, SD±7.50: education: 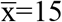 years, SD±2.68; sex 9F/31M) was included within 24 hours after injury upon meeting the American Congress of Rehabilitation Medicine (ACRM) criteria for mTBI (ACRM, 1993) in which the patient has to exhibit a traumatically induced physiological disruption of brain function as manifested by: i) any period of loss of consciousness (LOC), ii) any loss of memory for events immediately before or after the accident, iii) any alteration of mental state at the time of the accident (feeling dazed, disoriented, and/or confused), or iv) focal neurologic deficits that may or may not be permanent. Other inclusion criteria for this study were age between 18-55 years, acute brain CT as part of clinical care within 24 hours of injury, no significant polytrauma that would interfere with the follow-up and outcome assessment, no MRI contraindication. Fifteen patients reported history of anxiety or depression but none had a history of major psychiatric or neurological disorders. Visual acuity and hearing adequate for outcomes testing, fluency in English, and ability to give informed consent was required. Galveston Orientation and Amnesia Test (GOAT) score assessed at the time of informed consent was normal (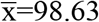; SD±2.18).

Fourteen orthopedic trauma control subjects matched by age (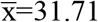 years; SD±10.14), education (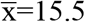 years; SD±2.17), and sex (6F/8M) were also recruited from the ED. Orthopedic injury causes included falls, pedestrian run overs, and bike accidents. All subjects presented with lower extremity fractures except for one that had an upper extremity fracture. Control subjects were ruled out for the current study if the emergency room physician required CT brain scan for suspicion of head trauma or if by interviewing medical services or subjects, participants reported clinical information such as loss of consciousness, amnesia, previous TBI, psychiatric or neurological prevalent pathology, and if, according to the abbreviated injury scale (AIS), this study would be counterproductive for their sustained systemic injuries.

All eligible subjects who voluntarily agreed to participate gave written informed consent. All study protocols were approved by the University of California, San Francisco Institutional Review Board.

### Neuropsychological Assessment

Commonly affected neuropsychological domains after mTBI were assessed using self-report and performance-based cognitive measures at 2 weeks and 6 month after injury: i) The Rivermead Postconcussion Symptoms Questionnaire (RPQ3-13), a self-reported questionnaire consisting of 16 physical and psychosocial symptoms frequently reported after mTBI; ii) the Rey Auditory Verbal Learning Test (RAVLT) to evaluate learning, short, and long-term memory; iii) Trail Making Tests A (TMTA) and B (TMTB) to evaluate attention, processing speed, and cognitive flexibility to switch tasks (TMTB-A); and iv) the Wechsler Adult Intelligence Scale (WAIS) coding and symbol search subscales for processing speed and visuo-perceptive association learning (Lezak et al., 2012). Scores reported for all measures were the raw scores.

### Image Acquisition

All mTBI subjects underwent a standardized MRI protocol acquired on a 3T GE MR750 scanner equipped with an eight channel phased array head radiofrequency coil (GE Healthcare, Waukesha, WI) at 2 weeks (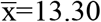 days; SD±2.10) and 6 months (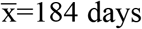 days; SD±8.86) after injury. Whole-brain DTI was performed with a multi-slice single-shot spin echo echoplanar pulse sequence (echo time [TE] = 81 ms; repetition time [TR] = 9 s) using 64 diffusion-encoding directions, isotropically distributed over the surface of a sphere with electrostatic repulsion, acquired at b = 1300 s/mm^2^ and another 64 directions at b = 3000 s/mm^2^, eight acquisitions at b = 0 s/mm^2^ for each set of 64 directions, slices of 2.7-mm thickness each with no gap between slices, a 128 x128 matrix, and a field of view (FOV) of 350 × 350 mm. Sagittal three-dimensional (3D) inversion recovery fast spoiled gradient recalled echo T1-weighted images (inversion time [TI] = 400 ms; flip angle, 11 degrees), were acquired with 256mm FOV, 200 contiguous partitions (1.2 mm) at 256×256 matrix. Sagittal 3D gradient echo T2*-weighted images (TE = 250 ms; TR = 500 ms; flip angle 10 degrees), were acquired with 256mm FOV, and 130 contiguous slices (1.6 mm) at 192×192 matrix. Sagittal 3D T2-weighted fluid-attenuated inversion recovery images (FLAIR; TE = 102 ms; TR = 575 ms; TI = 163 ms) were acquired with 256mm FOV, 184 contiguous slices (1.2 mm) at 256×256 matrix. The control group scans were acquired with the same parameters of acquisition as the mTBI patients, with data available for this study at the 2 week time point after orthopedic injury.

### MRI Image Processing and Analysis

An overview of the imaging analysis and statistical methods used in this study is presented in Figure 1.

**Figure 1.**
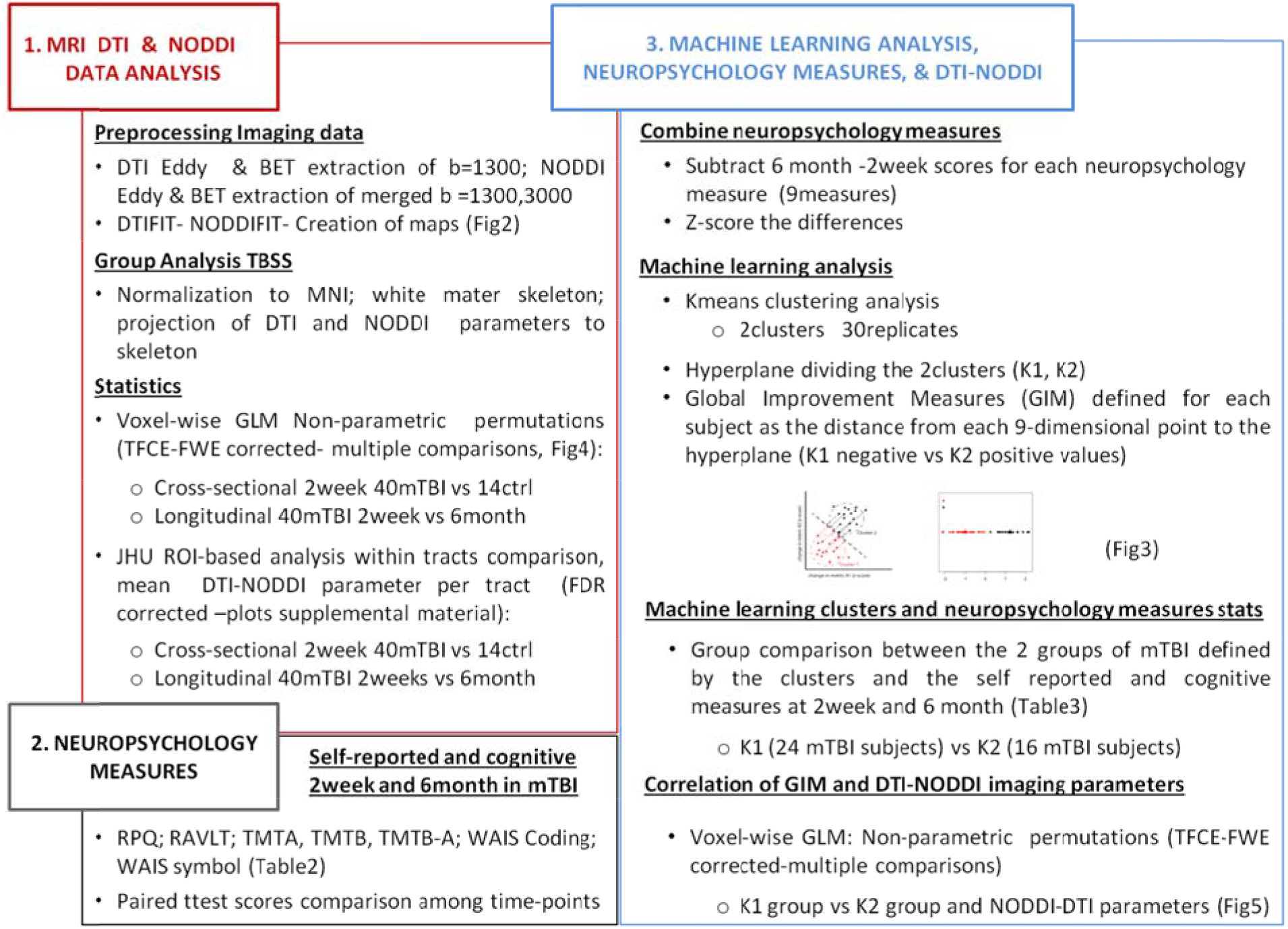
Overview of the imaging analysis and statistical methods.

### Radiological Findings

The structural 3T MRI images were interpreted by a board-certified neuroradiologist (E.L.Y), who was blinded to the initial presentation and subject group designation, using the NIH Common Data Elements (CDEs) for TBI pathoanatomic classification (Yuh et al., 2013). Supplementary Table 1 summarizes the clinical characteristics of the patients and the neuroradiological findings on structural 3T MRI. The MRI scans for the control group had no findings specific for TBI.

**Table 1.**
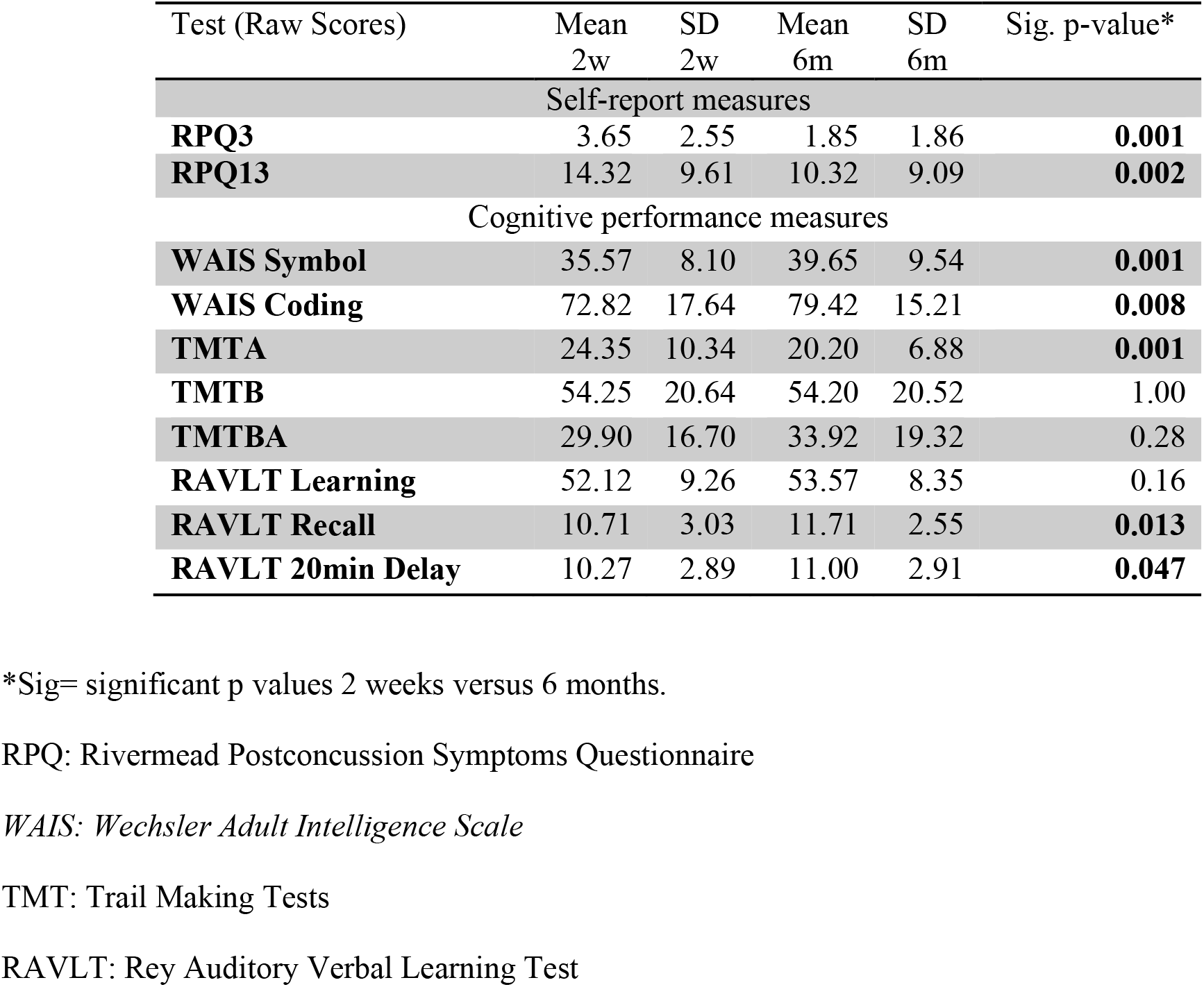
Self-Report and Cognitive-Performance Measures for the mTBI Patients

| Test (Raw Scores) | Mean<br>2w | SD<br>2w | Mean<br>6m | SD<br>6m | Sig. p-value* |
| --- | --- | --- | --- | --- | --- |
| Self-report measures |  |  |  |  |  |
| **RPQ3** | 3.65 | 2.55 | 1.85 | 1.86 | **0.001** |
| **RPQ13** | 14.32 | 9.61 | 10.32 | 9.09 | **0.002** |
| Cognitive performance measures |  |  |  |  |  |
| **WAIS Symbol** | 35.57 | 8.10 | 39.65 | 9.54 | **0.001** |
| **WAIS Coding** | 72.82 | 17.64 | 79.42 | 15.21 | **0.008** |
| **TMTA** | 24.35 | 10.34 | 20.20 | 6.88 | **0.001** |
| **TMTB** | 54.25 | 20.64 | 54.20 | 20.52 | 1.00 |
| **TMTBA** | 29.90 | 16.70 | 33.92 | 19.32 | 0.28 |
| **RAVLT Learning** | 52.12 | 9.26 | 53.57 | 8.35 | 0.16 |
| **RAVLT Recall** | 10.71 | 3.03 | 11.71 | 2.55 | **0.013** |
| **RAVLT 20min Delay** | 10.27 | 2.89 | 11.00 | 2.91 | **0.047** |

\*Sig= significant p values 2 weeks versus 6 months.
RPQ: Rivermead Postconcussion Symptoms Questionnaire
WAIS: Wechsler Adult Intelligence Scale
TMT: Trail Making Tests
RAVLT: Rey Auditory Verbal Learning Test

### Diffusion Tensor Imaging

The diffusion MRI data were verified to be free of major image artifacts or excessive patient movement, defined as more than 2mm of translation and/or rotation. DTI pre-processing and analysis were performed using tools from the Oxford Centre for Functional MRI of the Brain (FMRIB) Software Library, abbreviated as FSL (v. 5.0.7). First, images were corrected for eddy distortions and motion, using an average of the 8 b=0 s/mm^2^ volumes for each diffusion-weighted shell as a reference. The registered images were skull-stripped using the Brain Extraction Tool (Smith, 2002). All the resulting brain masks were visually inspected for anatomic fidelity. FA, MD, axial diffusivity (AD), and radial diffusivity (RD) maps were calculated using the FSL Diffusion Toolbox.

### Multi-compartment Biophysical Modeling of Diffusion MR Imaging

NODDI metrics were derived using the NODDI toolbox v0.9 (http://www.nitrc.org/projects/noddi_toolbox). We averaged the corresponding 8 b=0 s/mm^2^ images for each diffusion weighting. The NODDI code was modified to account for the slightly differing minimum echo times (TE) between images acquired at b = 1300 s/mm^2^ versus b = 3000 s/mm^2^ by fitting the NODDI model to the normalized diffusion-weighted images instead of the raw images. As per the developers’ recommendation, the diffusion-weighted images at each b value were normalized by the mean b = 0 s/mm^2^ images acquired with the same minimal TE scan parameter, generating images with TE-independent signal intensity, as we have described previously (Owen et al., 2014). NODDI fitting was performed with the NODDI Matlab Toolbox using the default settings (http://www.nitrc.org/projects/noddi_toolboxv0.9). Maps of NDI, ODI and FISO were generated. Figure 2 displays examples of DTI and NODDI map parameters.

### Statistics

#### Tract-Based Spatial Statistics

After calculation of the FA map, a voxel-wise statistical analysis of the FA data was performed using Tract-Based Spatial Statistics (TBSS). FA data were aligned into the common FMRIB58 FA template, which is in MNI152 (Montreal Neurological Institute) standard space, using the non-linear registration algorithm FNIRT (Jenkinson et al., 2002). Next, a mean FA image was created from the images for all the subject’s serial scans in this common space and thinned to generate a mean FA white matter skeleton that represented the center of all tracts common to the entire group of scans and thresholded at > 0.2. The aligned FA volume was then projected onto the skeleton by filling the skeleton with FA values from the nearest relevant tract center.

Twelve main fasciculi were studied using masks obtained from the Jülich histological atlas (JHU) mapped onto the standard MNI152 space and resampled to 1mm resolution. Binary mask images from the fasciculi of interest were used to mask the individual skeletonized maps previously registered to the MNI standard space using the non-linear tools in the TBSS procedure. Mean FA, MD, AD, RD, ODI, NDI, and FISO values were obtained from each subject’s white matter skeleton as well as each of the skeletonized regions of interest (ROIs). Right and left tracts were averaged into one single measurement.

We compared between subjects using a voxel-wise general linear model (GLM) analysis with permutation testing to correct for multiple comparisons (Nichols et al., 2002) using threshold-free cluster enhancement (TFCE), family-wise error corrected (FWE) at p≤0.05. An unpaired t-test was employed to compare cross-sectionally the group of patients and controls in the voxel-wise analysis at 2 weeks. A paired t-test was used to compare differences among DTI & NODDI measures within the patient group between 2 weeks and 6 months.

### Machine Learning Analysis

A percentage of mTBI patients remain functionally impaired after 1 year post-injury (McMahon et al., 2014). In an attempt to distinguish these patients in our cohort, we used unsupervised machine learning to derive a metric of cognitive and symptom improvement and link it to the imaging biomarkers. First, to obtain and define a global improvement measure (GIM) that best reflects their outcomes, we first subtracted the 2-week scores from the 6-month scores for each of the 9 self-reported and cognitive measures described in Table 1 for each subject, then used a Z-score transformation to normalize the values. Since each individual test could be noisy, we sought to combine them together into a single composite metric that we defined as the GIM. We approached this task through an unsupervised k-means clustering analysis with two clusters and thirty replicates in MATLAB 2012b (The MathWorks, Inc., Natick, Massachusetts, United States). We then calculated a hyperplane to separate these two clusters equidistantly and each subject’s GIM was defined as the signed (positive/negative) distance between the subject’s recovery status and this hyperplane. This distance can also be expressed as a weighted average of the various symptomatic and cognitive metrics (Fig 3a). Intuitively, this represents a data-driven method to combine the various self-reported and performance-based cognitive metrics to provide a wide degree of discrimination between patient outcome groups. We used two clusters because we were interested in distinguishing the patients with the best improvement from those who did not improve. While one cluster represents patients whose testing trend indicates overall improvement between two weeks and six months, the other cluster represents patients whose overall testing indicates a lack of, or in some cases a regression of, testing performance. Finally, we performed the voxel-wise comparison among clusters determined by the GIM measure with the DTI and NODDI metrics using randomise, the nonparametric permutation analysis tool in FSL, with TFCE correction for multiple comparisons at p<0.05.

**Figure 2.**
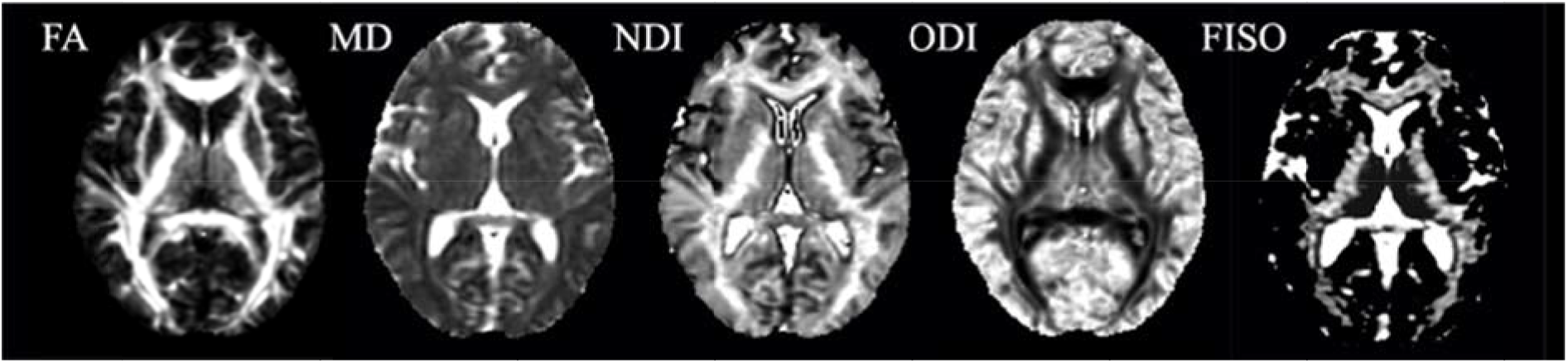
DTI and NODDI map examples. Example DTI (FA, MD) and NODDI (NDI, ODI, FISO) parameter maps obtained from a single subject in FMRIB58_FA standard-space.

**Figure 3.**
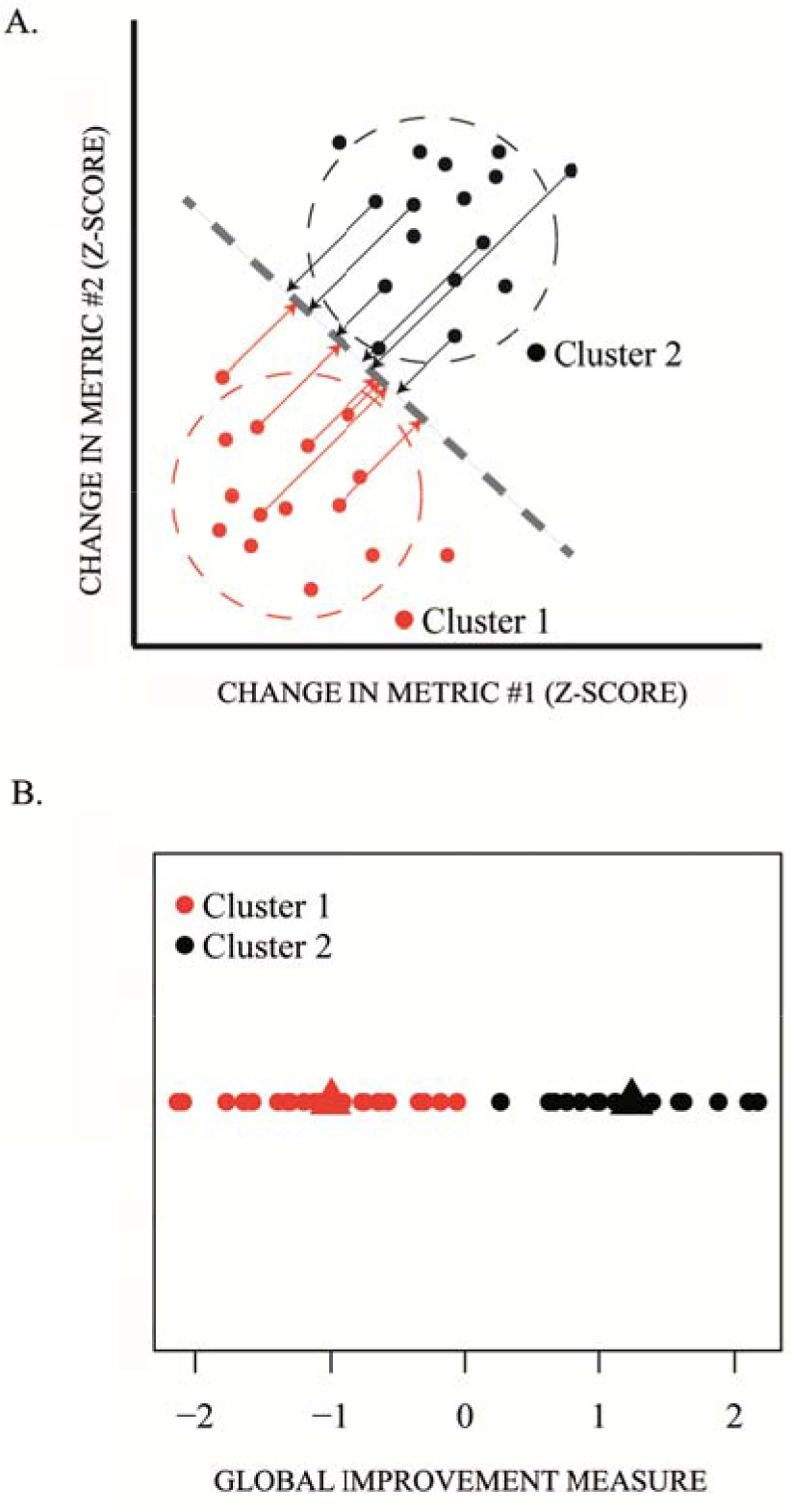
Machine Learning Clustering for Global Improvement Measure (GIM). (**A**): Two-dimensional schematic illustration of the clustering of change in cognitive and behavioral scores and the PCA projection used to define global improvement measure (arrows for some subjects demonstrating this distance). Subjects farther from the line in the upper right direction have better degree of recovery (black points), and subjects farther from the line in the other direction have less recovery (red points). (**B**): Two clusters comprising the actual group of mTBI patients based on their global improvement measure (GIM) showing Cluster 1 (red; n=24), patients with less recovery, and Cluster 2 (black; n=16), patients with more recovery. The triangles indicate the mean GIM for each cluster.

## RESULTS

### Neuropsychological Assessment

Table 1 displays the average results for the self-report and performance-based cognitive measures at 2 weeks and 6 months after injury for the 40 mTBI patients. Overall, patients self-reported a significant reduction in post-concussive symptoms on the RPQ scores, but a subset of subjects showed persistent self-reported symptomatology at the 6-month time point. Moreover, patients manifested improved performance in processing speed (TMTA), visuo-perceptive association learning (WAIS coding and symbol), as well as verbal memory (RAVLT) at 6 months vs 2 weeks.

### DTI and NODDI Voxel-wise Group Comparisons

#### Cross-sectional analysis between mild TBI patients and controls at the 2-week time point

Compared to the trauma controls, the mTBI patients showed decreased FA and increased MD in the genu and body of the corpus callosum, anterior and posterior limbs of the internal capsule, anterior corona radiata, anterior thalamic radiation, external capsule and cingulum. FISO was found to be increased in patients versus controls for the same tracts, but also additionally in the superior longitudinal fasciculi, posterior corona radiata, and fronto-occipital tracts. To a lesser extent, NDI also showed decreases mainly in the external capsule, anterior thalamic radiations, inferior longitudinal fasciculi, fornix and stria terminalis (Fig 4a). Figure 5 shows the mean values, effect sizes (Cohen’s d), and distributions of the sample for FA, MD, and FISO in the JHU tracts shown to be most affected by mTBI in the data-driven voxel-wise whole-brain TBSS analysis of Figure 4. The rest of the JHU tracts for the DTI and NODDI measures are provided in the Supplemental Material, Section A, Figs S1–S5.

**Figure 4.**
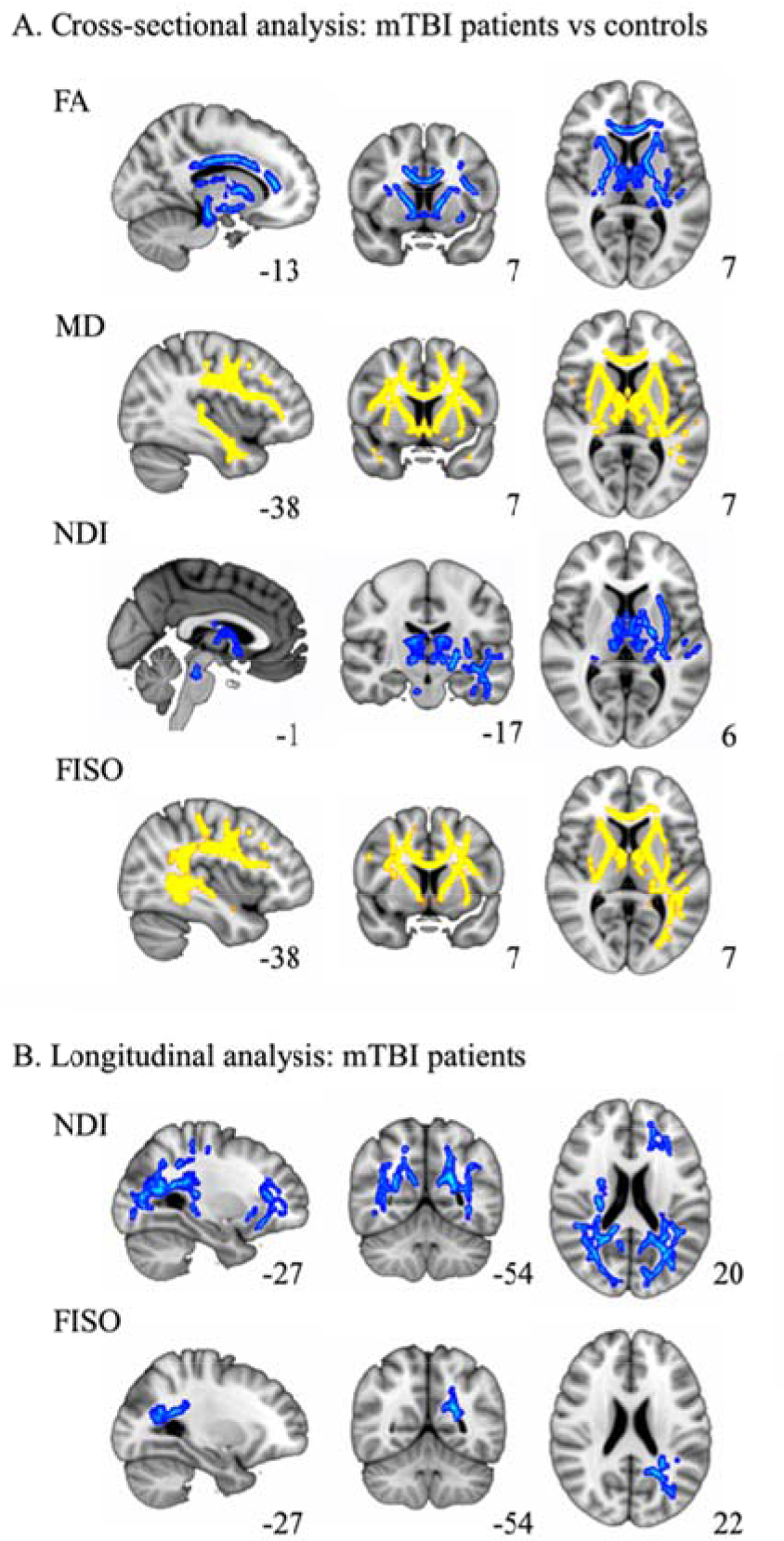
Cross-sectional and Longitudinal Voxel-wise Analysis. (**A**): Cross-sectional voxel-wise analysis comparison at 2 weeks between patients and controls. FA: fractional anisotropy; MD: mean diffusivity; ND: neurite dispersion index; FISO: volume fraction of isotropic water; In yellow: parameter increased in patients related to controls; In blue: parameter decreased in patients relative to controls. (**B**): Longitudinal voxel-wise comparison between 2 weeks and 6 months after injury in the patients group. In blue: parameter decreased over time. All results corrected for multiple comparisons using threshold-free cluster enhancement (TFCE) family-wise error (FWE) at p<0.05. The number next to each image is the MNI atlas coordinate defining its plane.

**Figure 5.**
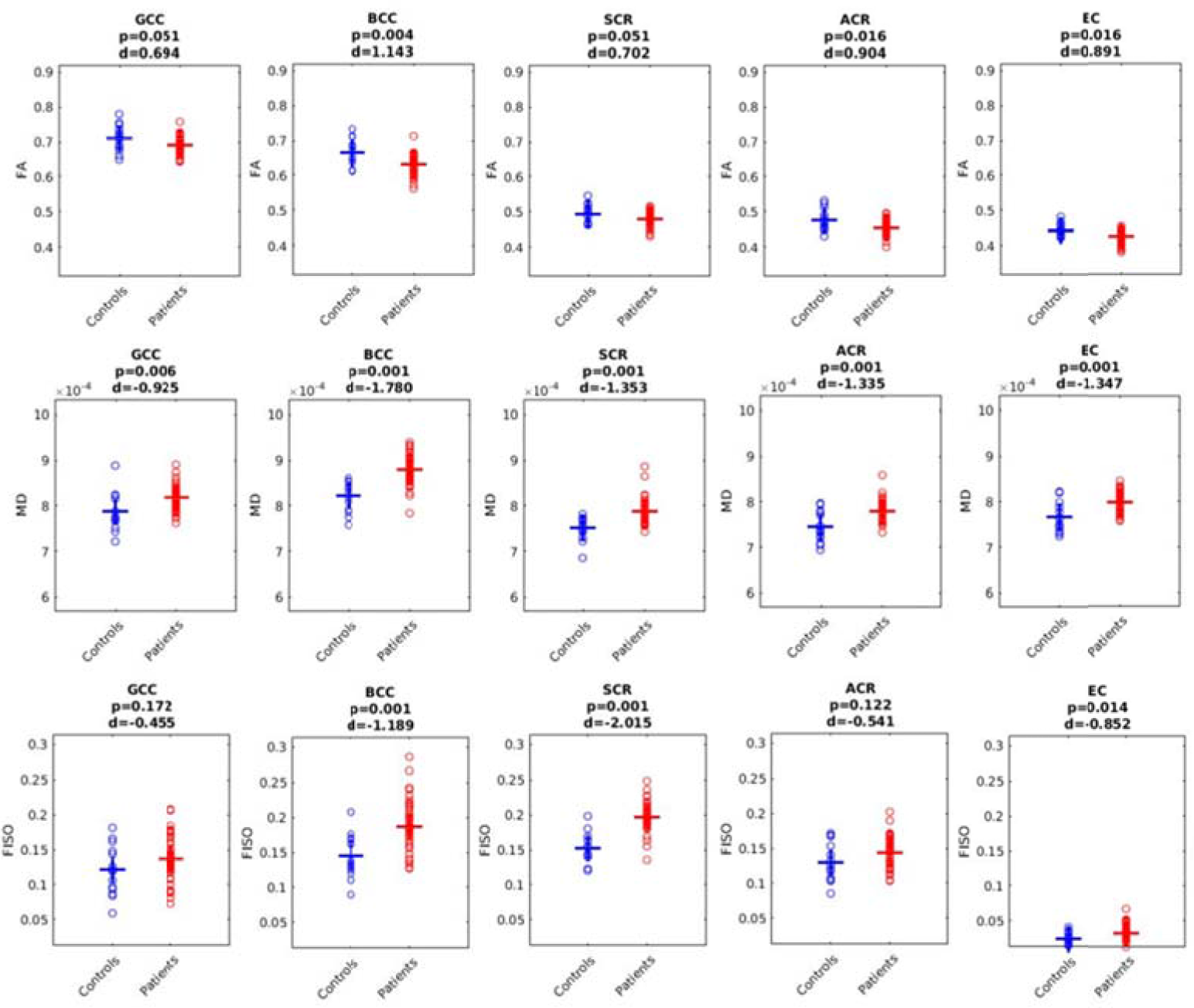
JHU Tracts Cross-sectional Comparison at 2 weeks Between mTBI and Controls. Averaged fractional anisotropy (FA), mean diffusivity (MD), and free water fraction (FISO) values of the left/right JHU tracts for each subject at the 2-week time point. GCC: genu corpus callosum; BCC: body corpus callosum; SCR: superior corona radiata; ACR: anterior corona radiata; EC: external capsule; Patient and control comparison FDR corrected p at 0.05. d: Cohen’s d effect size.

#### Longitudinal analysis of mTBI patients at 2 weeks vs 6 months

Longitudinal voxel-wise analysis of mTBI patients showed decreases over time of NDI in tracts including the anterior and posterior corona radiata, posterior thalamic radiation, inferior longitudinal and inferior fronto occipital fasciculi, posterior and anterior thalamic radiation, uncinate fasciculi, and external capsules. FISO showed decreases over time in the posterior corona radiata and inferior longitudinal fasciculi (Fig 4b). NODDI measures were more sensitive to microstructural damage in posterior tracts than DTI (Fig 6). Mean values and effect sizes, as well as the distribution of the samples, for all the DTI and NODDI measures are provided for the remaining JHU tracts in the Supplemental Material, Section B-Figs S6–S10. No other significant results were found for the rest of the DTI and NODDI parameters.

**Figure 6.**
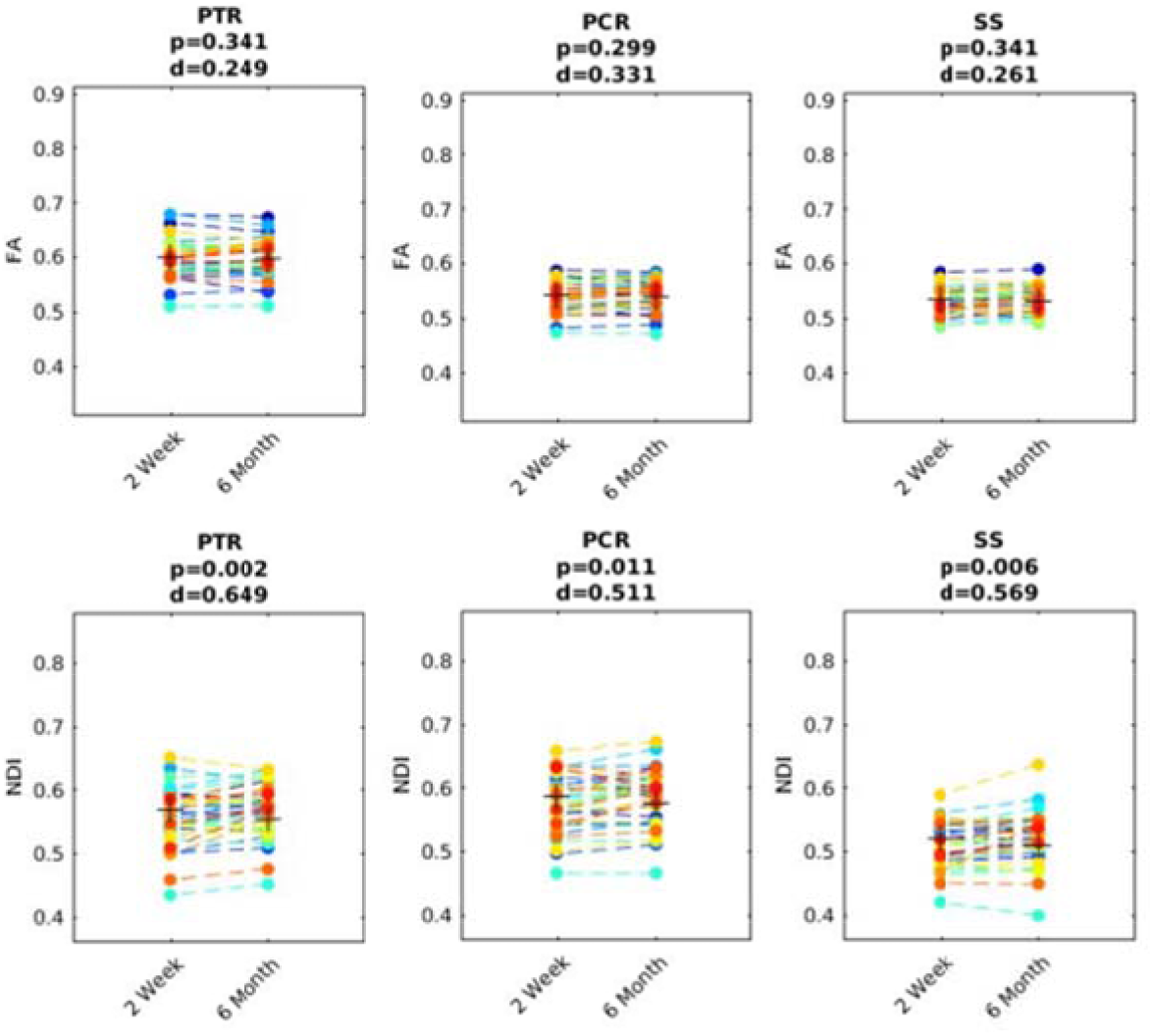
JHU Tracts Longitudinal Comparison at 2 weeks vs 6 month in mTBI. Averaged fractional anisotropy (FA) and neurite density index (NDI) values of the left/right JHU tracts for each subject at 2-week vs 6-month time points, showing significant changes over time in posterior tracts on NDI not captured by the FA parameter. PTR: posterior thalamic radiation; PCR: posterior corona radiata; SS: sagittal stratum (merged inferior fronto-occipital and inferior longitudinal fasciculi). Comparison FDR corrected p at 0.05. d: Cohen’s d effect size.

### Machine Learning Analysis

#### Machine learning clusters among mTBI patients and their global improvement measure

Two clear clusters (subgroups) were obtained dividing the group of mTBI patients based on their global improvement measure. Cluster K1 included 24 patients who had less improvement by the GIM metric than Cluster K2, which consisted of the 16 patients with the best global improvement (Fig 3b). Table 1 shows the change from 2 weeks to 6 months post-injury in the self-report and cognitive performance measures that comprise the GIM across all 40 mTBI patients. While the effect sizes of the change over time in the group means appear small for these measures, these group averages obscure variation among patients that can be uncovered by the unsupervised machine learning analysis dividing the group into two clusters based on the GIM. Is noteworthy to mention that, because of this division, K1 and K2 differed in years of education (K1: 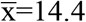 years, SD±2.1; K2: 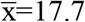 years; SD±2.5; p=0.002) but not in age.

#### Self-report and cognitive measures based on machine learning clustering

Supplementary Table 2 shows the difference between K1 and K2 in each of the nine measures that comprise the GIM. The patients of K2 were much more symptomatic on the RPQ than those of K1 at 2-weeks post-injury but recover to symptom levels similar to K1 by 6-months post-injury. The patients of K2 perform equivalently to those of K1 on the cognitive performance tests at 2-weeks post-injury, but, at the 6-month time point, are significantly outperforming their counterparts on the RAVLT and the WAIS, particularly the WAIS Coding subtest. K2 also trended toward better performance than K1 on the TMTA at both time points.

#### Voxel-wise group comparison of the DTI and NODDI parameters based on machine learning clustering

No significant relationship between traditional DTI metrics and patient global recovery GIM was found. However, NODDI metrics were found to be associated with cluster membership based on the GIM metric (Fig 7). The voxel-wise group comparison between K1 and K2 revealed increases in FISO in K2 compared to K1, but the pattern of elevated FISO varied between the 2-week and 6-month time points. The increased FISO of K2 versus K1 was posterior predominant at 2 weeks post-injury whereas the increased FISO of K2 versus K1 was anterior predominant at 6 months post-injury. In contradistinction, increased ODI of K1 versus K2 was found, both at 2 weeks and at 6 months in a largely stable pattern encompassing much of the central white matter tracts, with only the right internal capsule showing resolution of the elevated ODI at 6 months.

**Figure 7.**
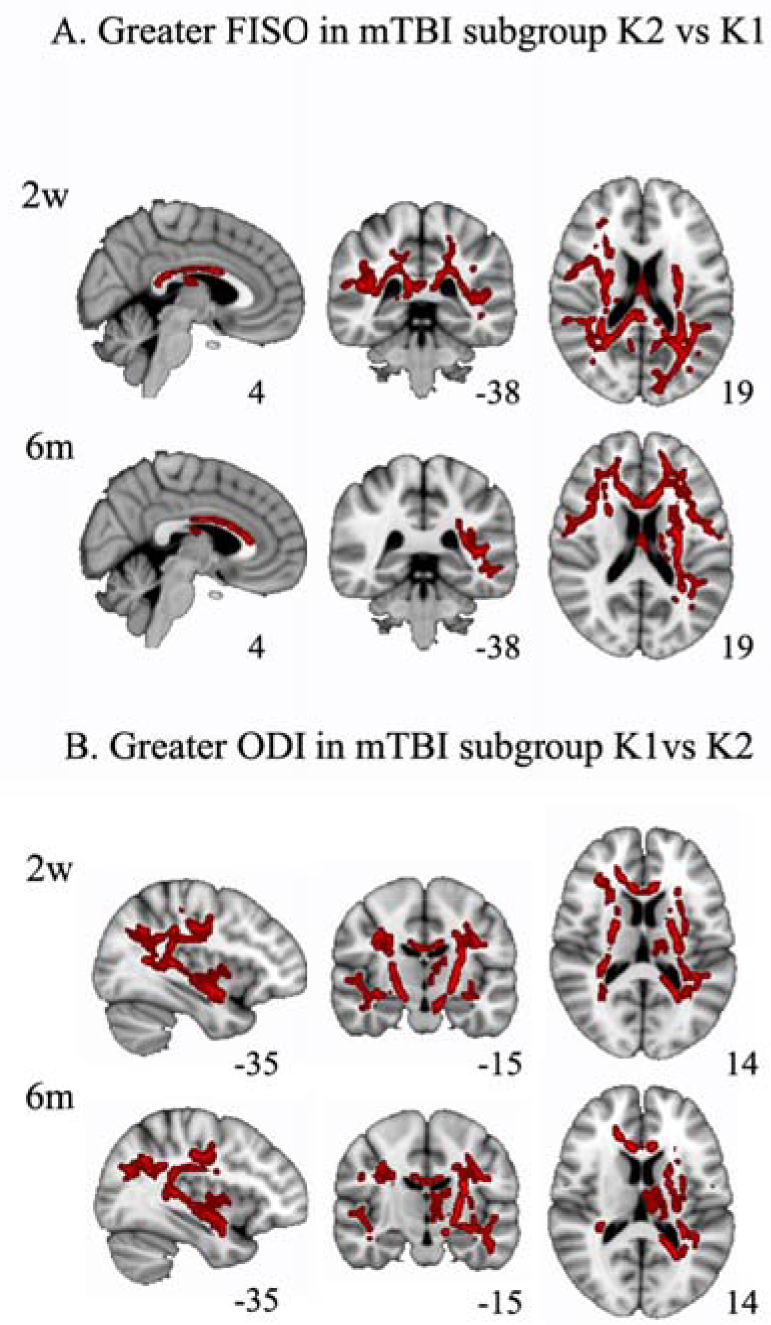
Voxel-wise Comparison of NODDI Metrics between mTBI Patient Subgroups K1 & K2. Voxel-wise group comparison of the DTI and NODDI parameters based on the machine learning cluster division (**A**): FISO: volume fraction of isotropic water, increased in patients in cluster 2 (K2). FISO increased in posterior tracts at 2 weeks while pattern at 6 months showed increases mainly in anterior tracts (**B**): ODI: orientation dispersion is increased in patients in cluster 1 (K1). All results corrected for multiple comparisons using threshold-free cluster enhancement (TFCE) family-wise error (FWE) at p<0.05. The number next to each image is the MNI atlas coordinate defining its plane.

## DISCUSSION

To the best of our knowledge, there is only one previous NODDI study of TBI, which was a cross-sectional investigation of sport-related concussion (Churchill et al., 2017). Our study compares DTI to NODDI longitudinally for the evolution of white matter microstructural injury and its association with symptomatic and cognitive recovery over time in a sample of civilian mTBI patients. The main findings are: i) early decreases in FA and NDI along with early increases of MD and FISO in the mTBI patient group versus trauma controls; ii) longitudinal white matter changes within the mTBI group as reflected by decreases in NDI and FISO over time; and iii) statically reduced ODI in those mTBI patients without symptomatic or cognitive improvement (K1) and dynamically elevated FISO in those mTBI patients with symptomatic recovery and progressively improved cognitive function (K2).

TBI involves multiple different time-varying pathophysiological effects, including diffuse axonal injury, diffuse microvascular injury, and neuroinflammation, that can lead to neurologic dysfunction (Bigler; Corps, JAMA). Due to this complexity, combining different biophysical measurements has potential for characterizing the underlying microarchitectural changes in the brain tissue (Cercignani et al., 2017). Our cross-sectional DTI findings at the 2-week time point showed decreases in FA and increases in MD in the mTBI group versus trauma controls, mainly in anterior tracts of the frontal and temporal lobes. These findings are in close agreement with prior DTI studies at this early stage of mTBI (Yuh et al., 2014; Eirud et al., 2014). In addition to these established DTI results, NODDI analysis revealed two additional findings at the 2-week time point. First, NDI was decreased in mTBI patients versus controls, primarily in the external capsule, thalamic radiations, inferior longitudinal fasciculi, and fornix- stria terminalis. Second, we also found increases in FISO in the same predominantly anterior distribution as FA and MD. Overall, these cross-sectional results suggest that the early decrease of FA and increase of MD after mTBI is due to an increase in free water content, possibly reflecting neuroinflammation. The sole prior study reporting DTI and NODDI measurements in TBI, in a sample of athletes with contact exposure, found increases in FA as well as increases in NDI and reduced ODI values (Churchill et al., 2017). These results may differ from those found in our study due to the mechanism of injury in young athletes involving repetitive subconcussive hits over long periods of time, thereby conflating injury and recovery effects, rather than a single episode of mTBI that can be serious enough to produce anatomic lesions on structural MRI (Supplementary Table 1).

Longitudinally within the group of mTBI patients, we only observed decreases over time in NDI values, suggesting progressive axonal degeneration, while there were no significant differences in the DTI parameters. This indicates that NDI is a more sensitive metric of white matter axonal loss than the conventional DTI metrics such as FA or MD. The decreases found in NDI over time were primarily in the bilateral posterior periventricular and left anterior periventricular white matter. It has been shown that a disproportionately high number of structural connectome links between gray matter areas traverse these regions of deep white matter. Indeed, these periventricular white matter regions have been described as a consistent nexus of network connectivity in the human brain (Owen et al., 2015; 2016). Additionally, virtual lesioning of these areas of deep periventricular white matter in tractography simulation experiments are particularly disruptive to the overall integrity of the whole-brain white matter network, demonstrating their singular importance to the large-scale structural connectome (Wang et al., 2017). Abnormal posterior periventricular white matter microstructure has also been described in sensory processing disorders (Owen 2013; Chang 2015) and were the only consistently affected white matter regions in a meta-analysis of DTI studies of attention-deficit hyperactivity disorder (ADHD, Chen et al., 2016). Furthermore, the global integrity of the structural connectome is linked to attention and executive function (Xiao et al., 2016). This constellation of recent results may account for the major impairments of concussions and mTBI, which are sensitivity to sensory stimuli (e.g., to bright lights and loud noises, as also seen in sensory processing disorder), attention deficits and executive dysfunction. Future studies combining microstructural characterization of these posterior periventricular white matter tracts with connectomic mapping may prove especially effective for explaining long-term symptomatic, cognitive and behavioral outcomes after mTBI.

Data-driven machine learning analysis of a composite global improvement measure based on nine symptom self-report and cognitive performance measures known to be affected in mTBI produced two patient clusters. One was a higher performing subgroup (K2) with early self-reported symptoms that resolved over time and who also improved in the information processing speed (WAIS Coding) and verbal memory (RAVLT) domains. The other was a lower functioning subgroup (K1) that displayed relatively few initial symptoms, but still performed less well than K2 on the cognitive tests, especially at the 6-month time point. Although no significant DTI differences were seen between these two mTBI subgroups, NODDI showed higher ODI throughout much of the central white matter in the K1 group at both time points. This greater fiber orientation dispersion in the low functioning K1 cluster may perhaps represent a premorbid characteristic influenced by their lower average educational level than K2, which might also imply lower cognitive reserve. The less well organized central white matter may help explain their poorer cognitive performance compared to the higher functioning K2 cluster. The difference in cognitive and educational levels between the subgroups might possibly also affect symptom reporting. Specifically, the greater number of symptoms reported by the K2 subgroup at the early time point may partly represent awareness of actual cognitive decline from baseline, followed by eventual return to baseline in both symptoms and cognition by 6m, at least at the subgroup level if not for every patient. In future studies with larger cohorts, separating function domains (i.e. self-reported measures from cognitive symptoms such as memory and processing speed) could be beneficial for more granular prediction of recovery (Levin et al., 2013).

Another finding using NODDI was the higher FISO of K2 versus K1 in predominantly posterior white matter at 2-weeks post-injury versus predominantly anterior white matter at 6-months post-injury. Since the overall group of 40 mTBI patients showed elevated anterior white matter free water, indicating vasogenic edema, at the early time point (Fig. 4a), this means that the K2 subgroup had more extensive early edema than K1 that also included posterior white matter. However, the anterior white matter edema resolved more slowly in K2 than K1, resulting in the relative elevation of free water in this distribution at the long-term time point. The greater extent of early white matter edema corresponds to the greater symptoms reported by the K2 subgroup at that time, with improving edema by 6 months post-injury matching their improvement in self-reported and cognitive performance measures. This observed association between white matter edema and the trajectory of symptomatic and cognitive recovery after mTBI requires more study to determine if there is a causal relationship. We do not discard other cofactors contributing to this symptomatology in addition to the trauma. Future studies should also investigate the effects of higher levels of stress and depression symptoms prior to and after the mTBI (Maurizio Bergamino, et al., 2016).

This study is a pilot longitudinal investigation of metrics from biophysical compartment modeling of diffusion MRI, exemplified by NODDI, for mTBI with comparison to standard DTI biomarkers. The orthopedic trauma group provides a rigorous control cohort that shares clinical and demographic features with the mTBI patients. The results provide support for our original hypotheses of elevated free water fraction early after injury and of serial decline in white matter axonal density during the first 6 months post-trauma. Exploratory machine learning analysis of the relationship between NODDI measures and symptomatic/cognitive recovery show that (i) dynamically increased free water fraction across semi-acute and chronic time points was associated with better recovery, suggesting a beneficial role for edema/neuroinflammation; and (ii) statically reduced fiber orientation dispersion was correlated with better long-term cognition, consistent with prior studies showing more highly organized white matter in those with better intellectual functioning in multiple domains (Fijell, et al. 2011; Yang Y et al., 2015; Roger A Kievit, et al., 2016). These new hypotheses from the exploratory findings need to be tested in larger cohorts that have better statistical power for determining imaging-cognition relationships. In the absence of cognitive control data, we also cannot exclude a learning component for the improvement across the two cognitive assessments, especially for the better recovery subgroup that had, on average, a higher educational level than the poorer recovery subgroup. Another limitation of the study is the small sample size of the controls, which lack a longitudinal component.

In summary, we found that NODDI parameters are sensitive imaging biomarkers for the subtle yet complex underlying white matter microstructural pathology after mTBI, such as diffuse axonal injury and neuroinflammation. Our results show that the early decrease of FA and increase of MD after mTBI, which are primarily in the anterior white matter, correspond to white matter regions of elevated FISO, which may reflect inflammatory vasogenic edema. This elevation of free water is more extensive in the subgroup of patients reporting more post-concussive symptoms early after trauma. The longer-term changes from 2 weeks to 6 months after mTBI are marked by declining neurite density in predominantly posterior white matter, suggesting axonal degeneration from DAI for which NODDI appears more sensitive than any of the DTI metrics, such as FA. The affected posterior white matter regions are known to be topologically integral to the structural connectome and are involved in multiple sensory and cognitive domains, including attention and executive function. The observation of stably elevated white matter fiber orientation dispersion in the mTBI subgroup with poorer cognitive performance may represent the sensitivity of ODI to premorbid intellectual functioning. Further research studies in larger well-phenotyped cohorts are needed to validate these NODDI biomarkers for mTBI diagnosis, for prediction of symptoms and cognitive performance, and for treatment monitoring.

## Supplementary material

**Table 1.**
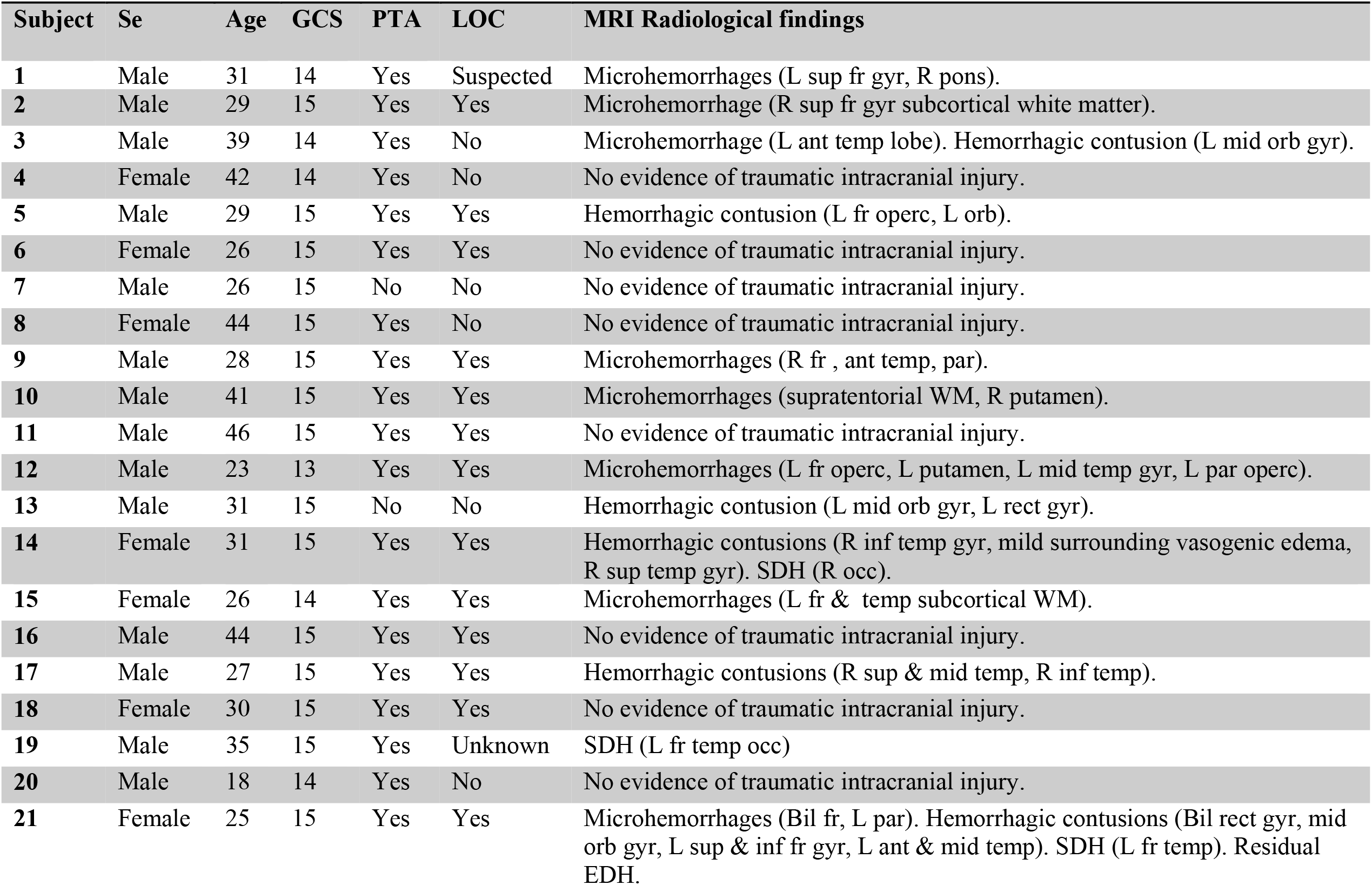
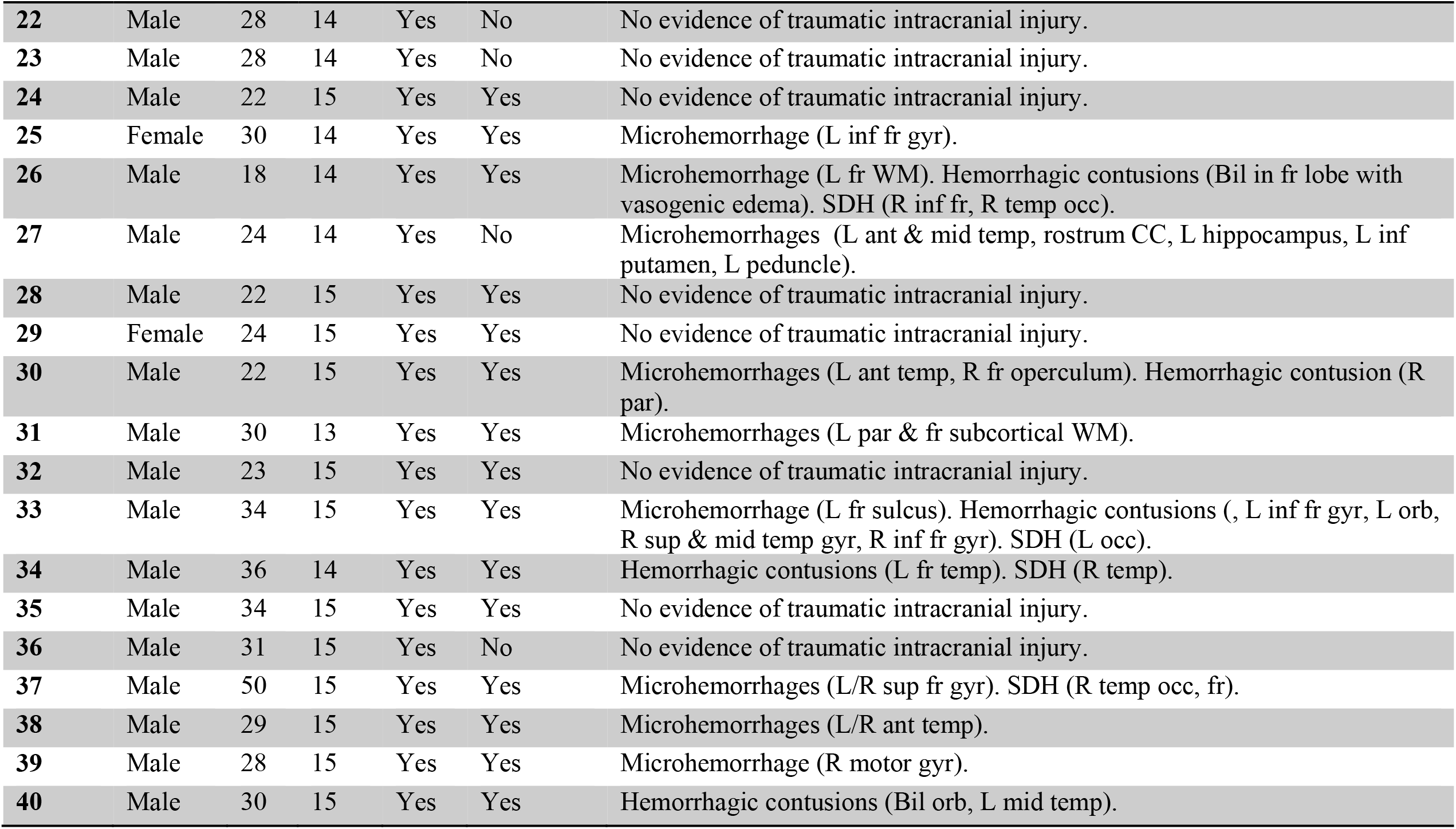
Demographic and clinical characteristics of mTBI patients

**Table 2.** Self-report and cognitive-performance measures per cluster based on mTBI recovery: K1 (worse improvement) & K2 (better improvement).

| Group Cluster Statistics 2 weeks K1 & K2 |  |  |  |  |
| --- | --- | --- | --- | --- |
| Test | Cluster group(K) | N | Mean | p-value* |
| RPQ3 | 1 | 24 | 2.37 | .000 |
|  | 2 | 16 | 5.56 |  |
| RPQ13 | 1 | 24 | 10.91 | .005 |
|  | 2 | 16 | 19.43 |  |
| WAIS Symbol | 1 | 24 | 35.58 | .994 |
|  | 2 | 16 | 35.56 | .994 |
| WAIS Coding | 1 | 24 | 72.29 | .818 |
|  | 2 | 16 | 73.62 |  |
| TMTA | 1 | 24 | 26.58 | .095 |
|  | 2 | 16 | 21.00 |  |
| TMTB | 1 | 24 | 56.41 | .423 |
|  | 2 | 16 | 51.00 |  |
| TMTBA | 1 | 24 | 29.83 | .976 |
|  | 2 | 16 | 30.00 |  |
| RAVLT Learning | 1 | 24 | 50.08 | .088 |
|  | 2 | 16 | 55.18 |  |
| RAVLT 20min Delay | 1 | 24 | 9.91 | .345 |
|  | 2 | 16 | 10.81 |  |

| Group Statistics 6months K1& K2 |  |  |  |  |
| --- | --- | --- | --- | --- |
| Test | Cluster group (K) | N | Mean | p-value* |
| RPQ3 | 1 | 24 | 1.95 | .658 |
|  | 2 | 16 | 1.68 |  |
| **RPQ13** | 1 | 24 | 10.41 | .939 |
|  | 2 | 16 | 10.18 |  |
| **WAIS Symbol** | 1 | 24 | 36.50 | .127 |
|  | 2 | 16 | 44.37 |  |
| **WAIS Coding** | 1 | 24 | 76.41 | **.009** |
|  | 2 | 16 | 83.93 |  |
| **TMTA** | 1 | 24 | 21.79 | .073 |
|  | 2 | 16 | 17.81 |  |
| **TMTB** | 1 | 24 | 55.75 | .578 |
|  | 2 | 16 | 52.00 |  |
| **TMTBA** | 1 | 24 | 33.75 | .945 |
|  | 2 | 16 | 34.18 |  |
| **RAVLT Learning** | 1 | 24 | 50.87 | **.010** |
|  | 2 | 16 | 57.62 |  |
| **RAVLT 20min Delay** | 1 | 24 | 9.91 | **.003** |
|  | 2 | 16 | 12.62 |  |

\*Bold p-values are significant at $p < 0.05$ , FDR corrected.
RPQ: Rivermead Postconcussion Symptoms Questionnaire
**WAIS: Wechsler Adult Intelligence Scale**
TMT: Trail Making Tests
RAVLT: Rey Auditory Verbal Learning Test.

### A. CROSS-SECTIONAL DTI & NODDI PARAMETERS: 40 mTBI patients vs 14 controls group comparison @ 2 weeks

**Glossary**
GCC: genu corpus callosum
BCC: body corpus callosum
SCC: splenium corpus callosum
ALIC: anterior limb of internal capsule
PLIC: posterior limb of internal capsule
ACR: anterior corona radiata
SCR: superior corona radiata
PTR: posterior thalamic radiation
EC: external capsule
CGC: cingulum
SLF: superior longitudinal fasciculi
SFO: superior fronto-occipital fasciculi
PCR: posterior corona radiate
SS: sagittal stratum (merged inferior fronto-occipital and inferior longitudinal fasciculi)

**Fig. S1.**
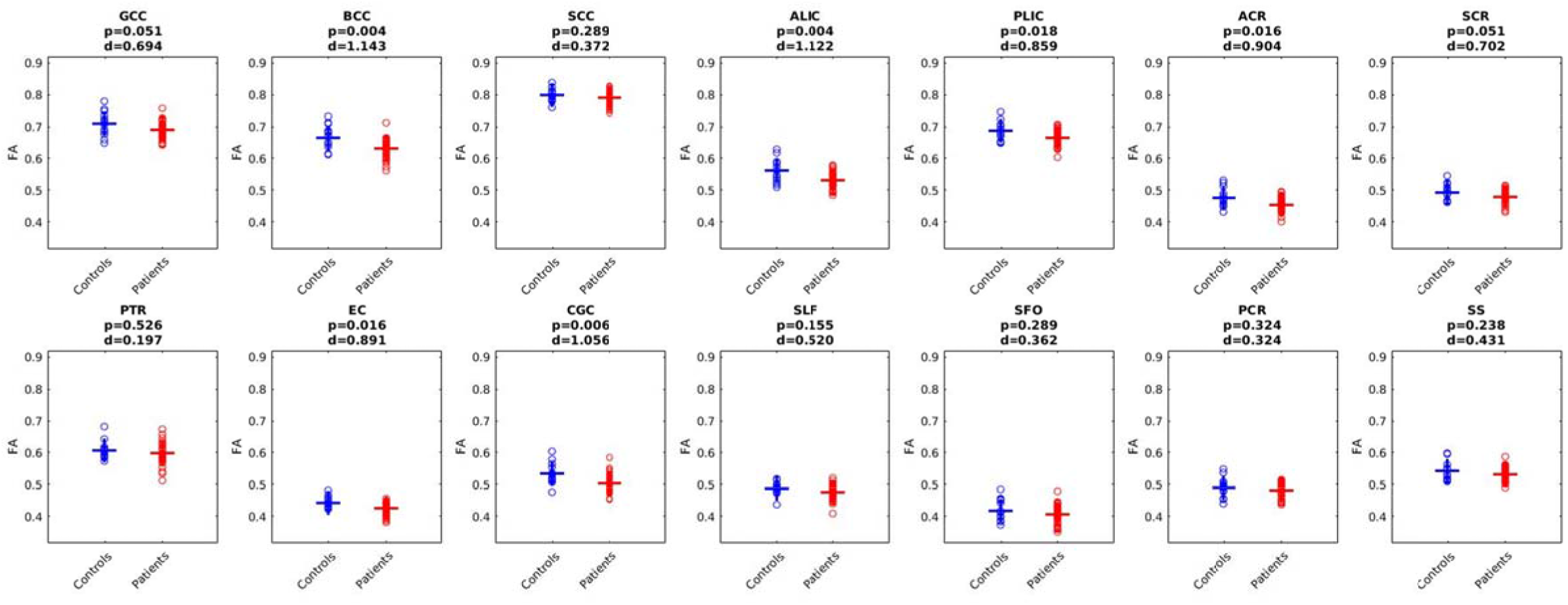
Averaged fractional anisotropy (FA) per left/right JHU tracts for each subject. FDR corrected p at 0.05; d: Cohen’s d effect size.

**Fig. S2.**
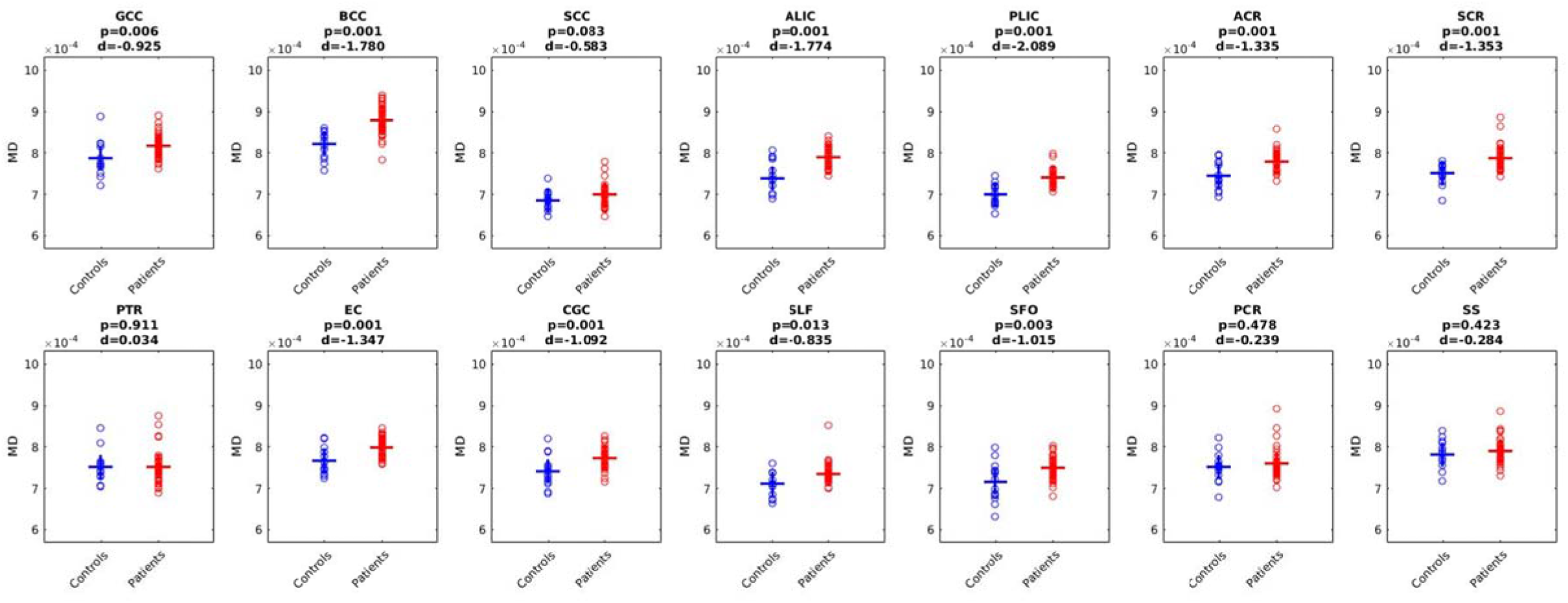
Mean diffusivity (MD) per left/right JHU tracts for each subject. FDR corrected p at 0.05; d: Cohen’s d effect size.

**Fig. S3.**
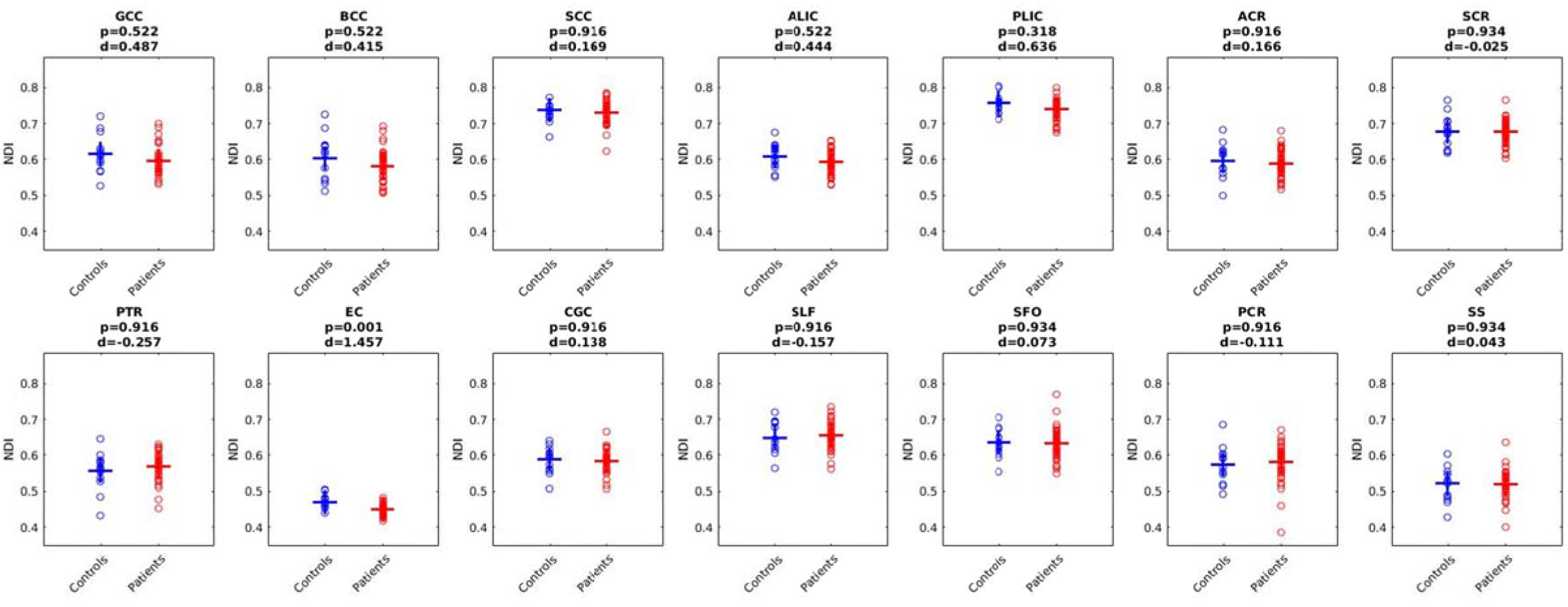
Averaged neurite density index (NDI) per left/right JHU tracts for each subject. FDR corrected p at 0.05; d: Cohen’s d effect size.

**Fig. S4.**
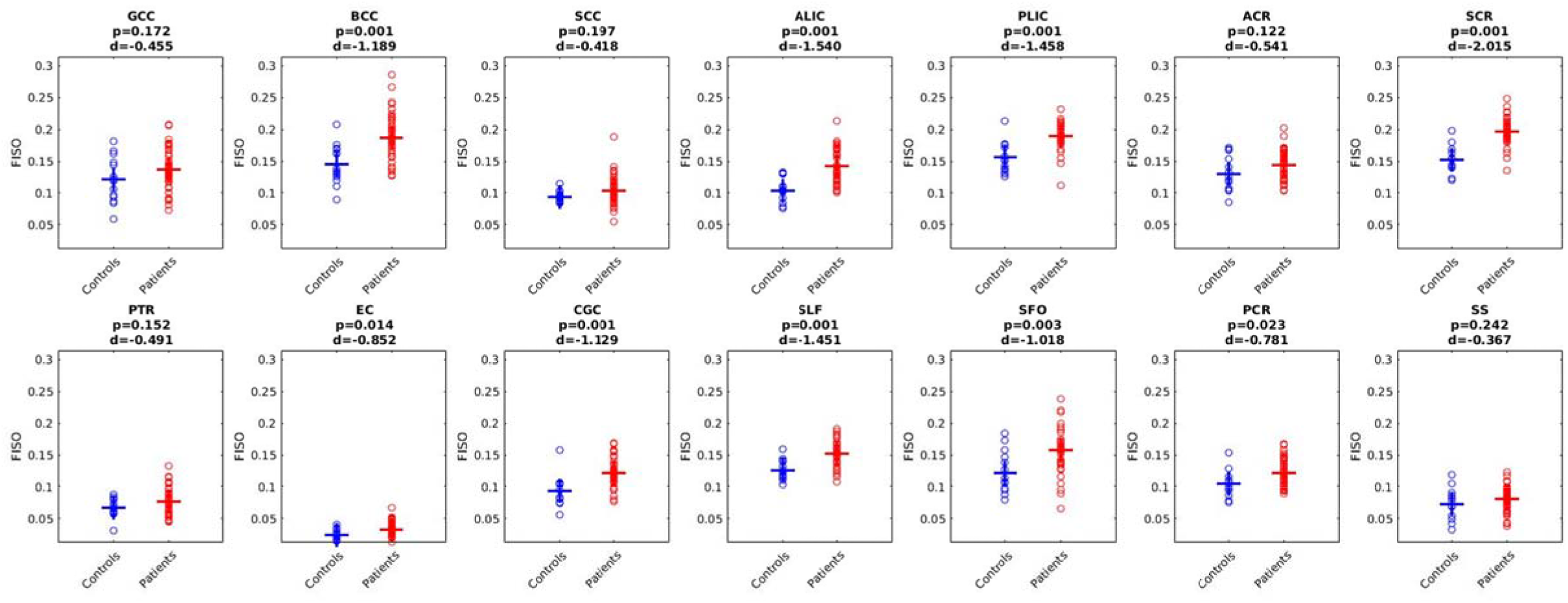
Averaged free water fraction (FISO) per left/right JHU tracts for each subject. FDR corrected p at 0.05; d: Cohen’s d effect size.

**Fig. S5.**
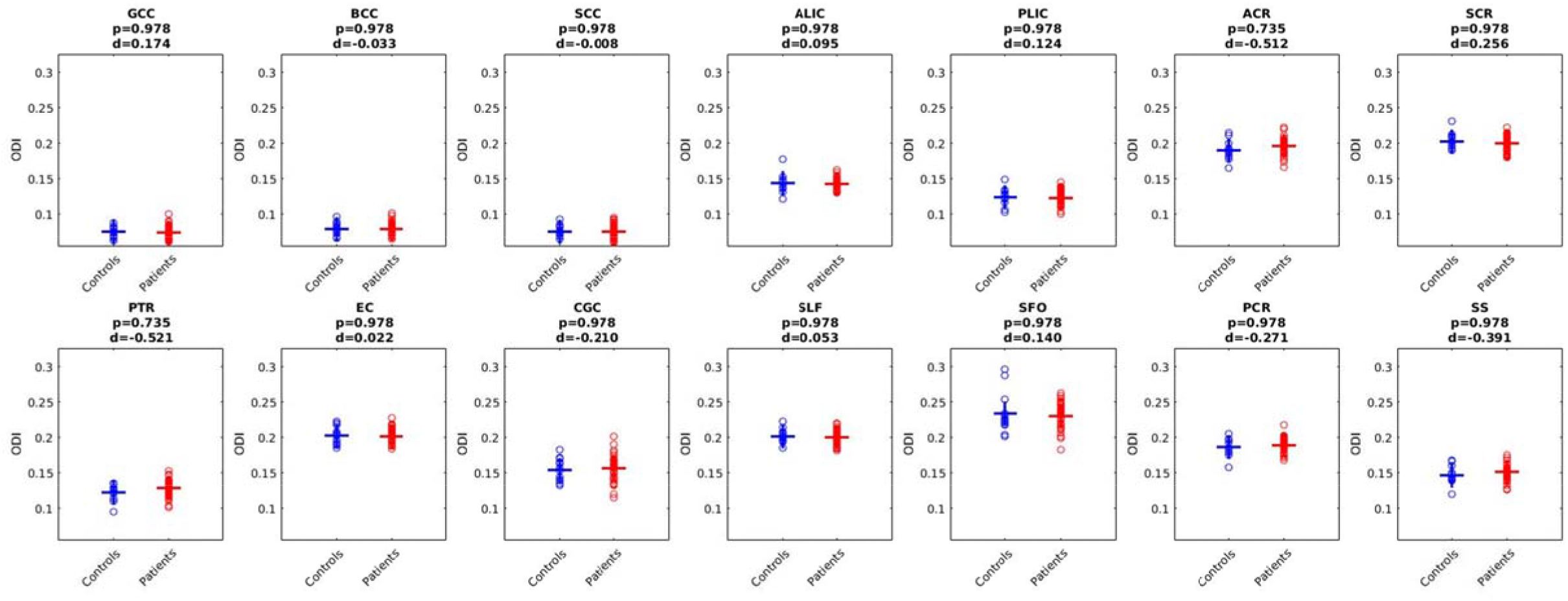
Averaged orientation dispersion (ODI) per left/right JHU tracts for each subject. FDR corrected p at 0.05; d: Cohen’s d effect size.

### B. LONGITUDINAL DTI & NODDI PARAMETERS: 40mTBI patients @ 2weeks vs 6month

**Fig. S6.**
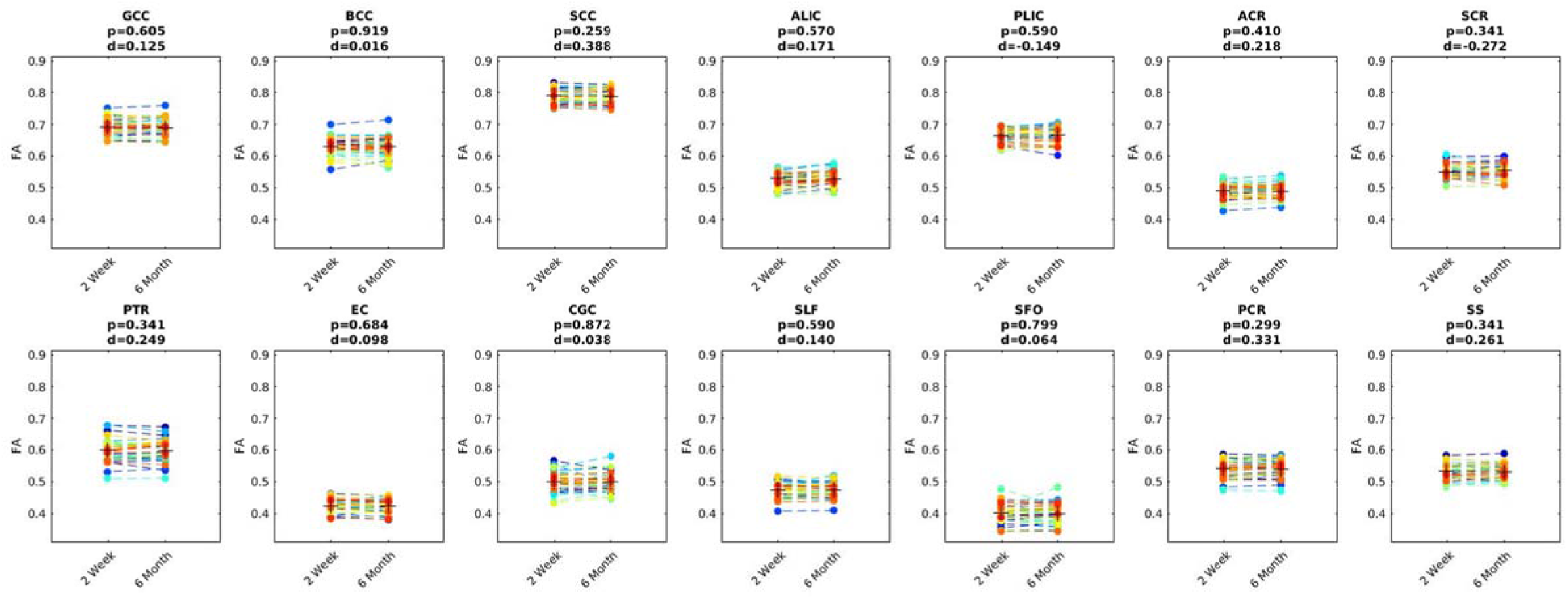
Averaged fractional anisotropy (FA) left/right JHU tracts for each subject at 2 week and 6 month time point. FDR corrected p at 0.05; d: Cohen’s d effect size.

**Fig. S7.**
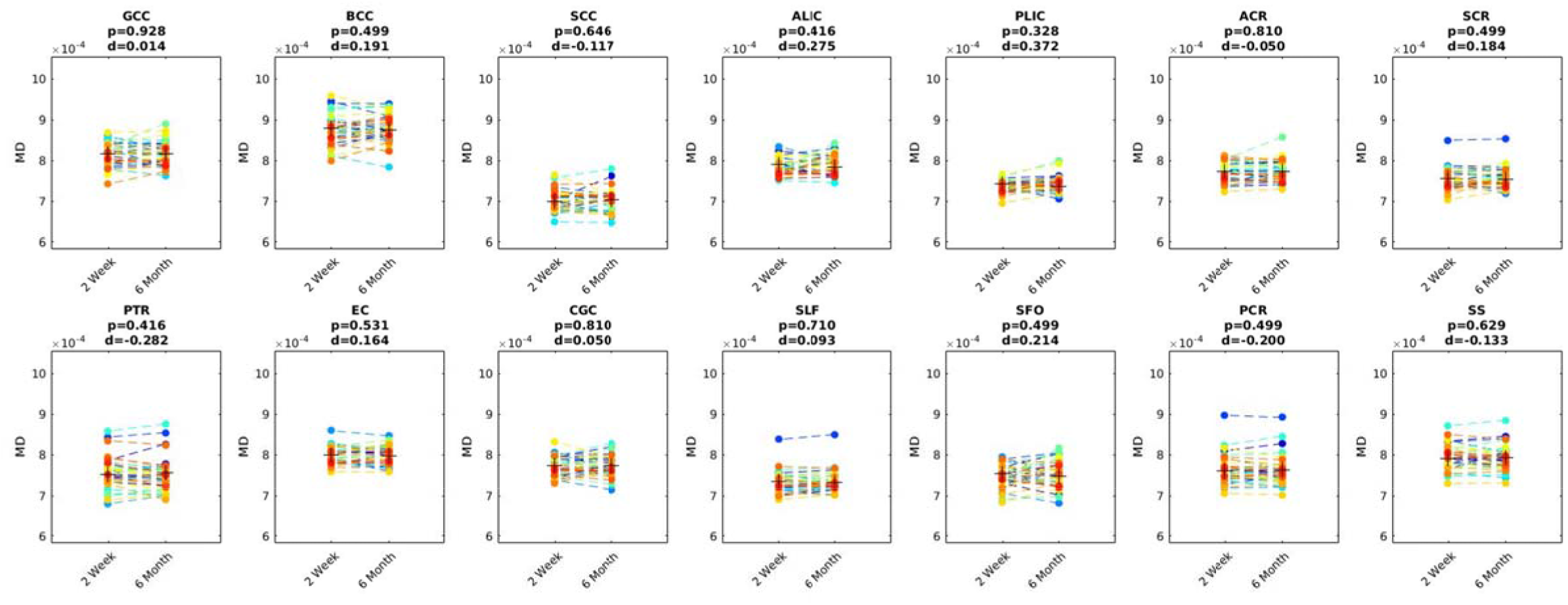
Averaged mean diffusivity (MD) left/right JHU tracts for each subject at 2 week and 6 month time point. FDR corrected p at 0.05; d: Cohen’s d effect size.

**Fig. S8.**
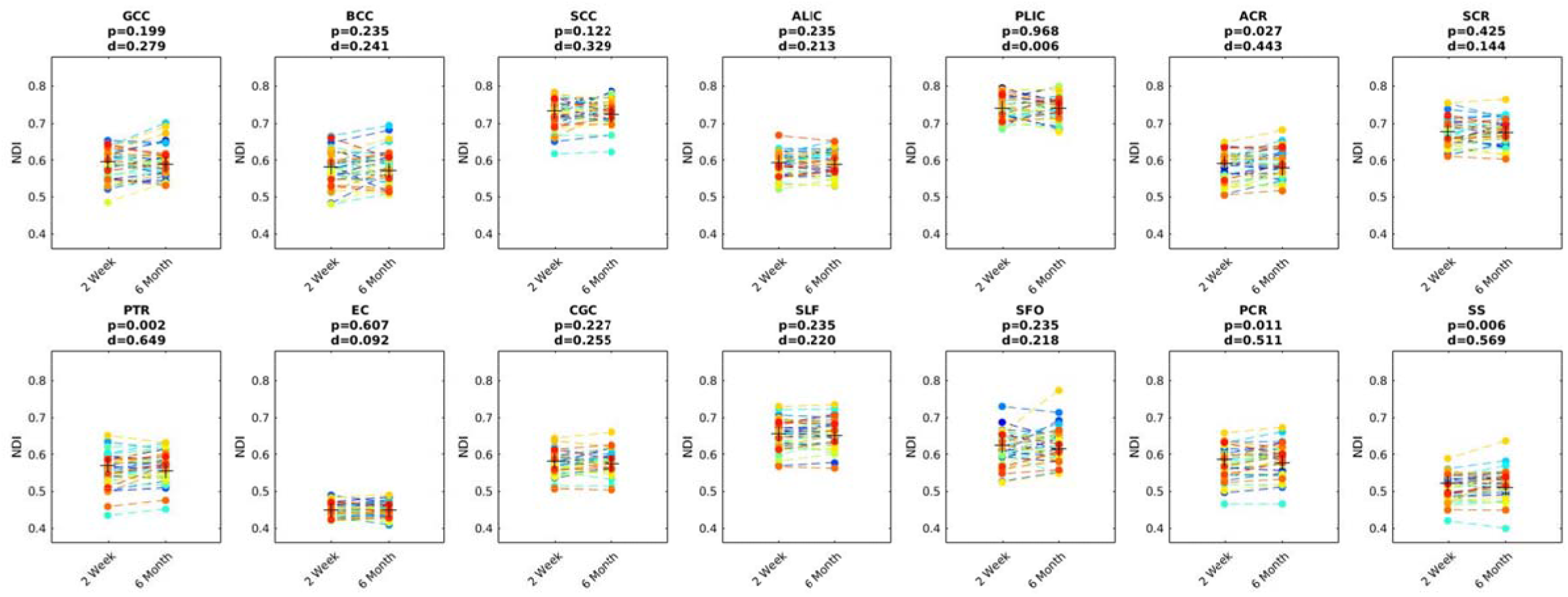
Averaged neurite density (NDI) left/right JHU tracts for each subject at 2 week and 6 month time point. FDR corrected p at 0.05; d: Cohen’s d effect size.

**Fig. S9.**
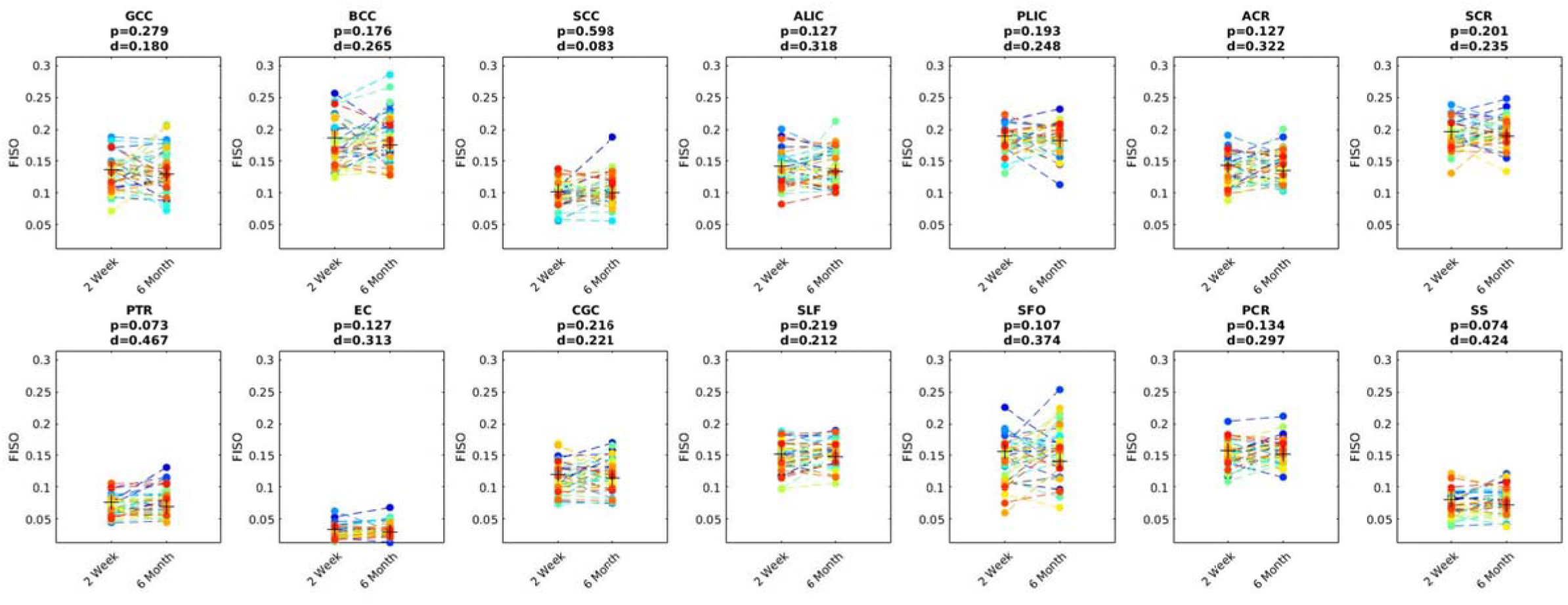
Averaged free water fraction (FISO) left/right JHU tracts for each subject at 2 week and 6 month time point. FDR corrected p at 0.05; d: Cohen’s d effect size.

**Fig. S10.**
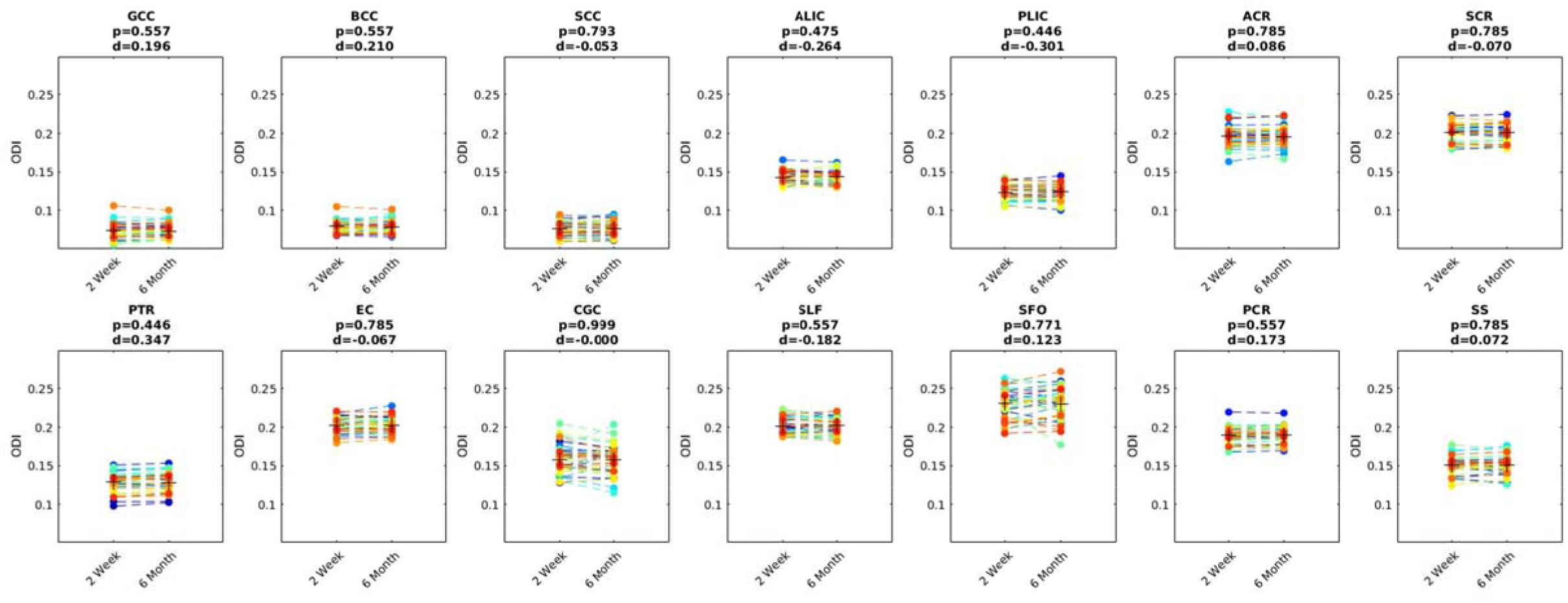
Averaged orientation dispersion (ODI) left/right JHU tracts for each subject at 2 week and 6 month time point. FDR corrected p at 0.05; d: Cohen’s d effect size.

## REFERENCES

1. American Congress of Rehabilitation Medicine (ACRM). Mild Traumatic Brain Injury Committee. (1993). Definition of mild traumatic brain injury. Journal of Head Trauma Rehabilitation 8 (3), 86–87.

2. Basser PJ, Jones DK. Diffusion-tensor MRI: theory, experimental design and data analysis - a technical review. NMR Biomed 2002;15:456–67. Review.

3. Bergamino M, Pasternak O, Farmer M, Shenton ME, Hamilton JP. Applying a free-water correction to diffusion imaging data uncovers stress-related neural pathology in depression. Neuroimage Clin 2015; 30:336–42.

4. Bigler ED. Neuroimaging biomarkers in mild traumatic brain injury (mTBI). Neuropsychol Rev 2013;23:169–209.

5. Bigler ED. Neuropathology of Mild Traumatic Brain Injury: Correlation to Neurocognitive and Neurobehavioral Findings. In: Kobeissy FH, editor. Brain Neurotrauma: Molecular, Neuropsychological, and Rehabilitation Aspects. Boca Raton (FL): CRC Press/Taylor & Francis; 2015. Chapter 31.

6. Billiet T, Vandenbulcke M, Mädler B, Peeters R, Dhollander T, Zhang H, Deprez S, Van den Bergh BR, Sunaert S, Emsell L. Age-related microstructural differences quantified using myelin water imaging and advanced diffusion MRI. Neurobio Aging 2015;36:2107–21.

7. Caverzasi E, Papinutto N, Castellano A, Zhu AH, Scifo P, Riva M, Bello L, Falini A, Bharatha A, Henry RG. Neurite Orientation Dispersion and Density Imaging Color Maps to Characterize Brain Diffusion in Neurologic Disorders. J Neuroimaging 2016;26:494–8.

8. Cercignani M, Bouyagoub S. Brain microstructure by multi-modal MRI: Is the whole greater than the sum of its parts? Neuroimage. 2017. pii: S1053-8119(17)30886-8.

9. Chang YS, Owen JP, Pojman NJ, Thieu T, Bukshpun P, Wakahiro ML, Berman JI, Roberts TP, Nagarajan SS, Sherr EH, Mukherjee P. White Matter Changes of Neurite Density and Fiber Orientation Dispersion during Human Brain Maturation. PLoS One 2015;10:e0123656.

10. Chen L, Hu X, Ouyang L, He N, Liao Y, Liu Q, Zhou M, Wu M, Huang X, Gong Q. A systematic review and meta-analysis of tract-based spatial statistics studies regarding attention-deficit/hyperactivity disorder. Neuroscience and biobehavioral reviews 2016;68:838–47.

11. Churchill NW, Caverzasi E, Graham SJ, Hutchison MG, Schweizer TA. White matter microstructure in athletes with a history of concussion: Comparing diffusion tensor imaging (DTI) and neurite orientation dispersion and density imaging (NODDI). Hum Brain Mapp 2017;38:4201–4211.

12. Croall ID, Cowie CJ, He J, Peel A, Wood J, Aribisala BS, Mitchell P, Mendelow AD, Smith FE, Millar D, Kelly T, Blamire AM. White matter correlates of cognitive dysfunction after mild traumatic brain injury. Neurology 2014;83:494–501.

13. Eierud C, Craddock RC, Fletcher S, Aulakh M, King-Casas B, Kuehl D, LaConte SM. Neuroimaging after mild traumatic brain injury: review and meta-analysis. Neuroimage Clin 2014;4:283–94.

14. Fijell AM, Westlye LT, Amlien IK, Walhovd KB. Reduced white matter integrity is related to cognitive instability. J Neurosci 2011; 31:18060–72.

15. Jelescu OI, Budde MD. Design and validation of diffusion MRI models of white matter. Frontiers in Physics 2017; 5:61. Review.

16. Jenkinson M., Bannister P, Brady M, Smith S. Improved optimization for the robust and accurate linear registration and motion correction of brain images. Neuroimage 2002;17: 825–41.

17. Jones DK, Cercignani M. Twenty-five pitfalls in the analysis of diffusion MRI data. NMR Biomed 2010; (7):803–20. Review.

18. Kamagata K, Zalesky A, Hatano T, Ueda R, Di Biase MA, Okuzumi A, Shimoji K, Hori M, Caeyenberghs K, Pantelis C, Hattori N, Aoki S. Gray Matter Abnormalities in Idiopathic Parkinson’s Disease: Evaluation by Diffusional Kurtosis Imaging and Neurite Orientation Dispersion and Density Imaging. Hum Brain Mapp 2017; 38: 3704–3722.

19. Kievit RA, Davis SW, Griffiths J, Correia MM, Cam-Can, Henson RN. A watershed model of individual differences in fluid intelligence. Neuropsychologia 2016; 186-106:417.

20. Levin HS, Diaz-Arrastia RR. Diagnosis, prognosis, and clinical management of mild traumatic brain injury. Lancet Neurol 2015;14:506–17.

21. Levin HS, Li X, McCauley SR, Hanten G, Wilde EA, Swank P. Neuropsychological outcome of mTBI: a principal component analysis approach. J Neurotrauma 2013;30:625–32.

22. Lezak, M D, Howieson, D B, Bigler, E D, & Tranel, D (2012). Neuropsychological assessment (5^th^ ed.). New York: Oxford University Press.

23. Mah A, Geeraert B, Lebel C. Detailing neuroanatomical development in late childhood and early adolescence using NODDI. PLoS One 2017;12:e0182340.

24. Manley GT, Maas AI. Traumatic brain injury: an international knowledge-based approach. JAMA 2013;310:473–4.

25. McMahon PJ, Hricik A, Yue JK, et al. Symptomatology and functional outcome in mild traumatic brain injury: results from the prospective TRACK-TBI study. Journal of neurotrauma 2014;31:26–33.

26. Mori S, Zhang J. Principles of diffusion tensor imaging and its applications to basic neuroscience research. Neuron 2006; 51:527–39. Review.

27. Mukherjee P, Berman JI, Chung SW, Hess CP, Henry RG. Diffusion tensor MR imaging and fiber tractography: theoretic underpinnings. AJNR Am J Neuroradiol 2008;29:632–41.

28. Nichols TE, Holmes AP. Nonparametric permutation tests for functional neuroimaging: a primer with examples. Hum Brain Mapp 2002; 15: 1–25.

29. Oehr L, Anderson J. Diffusion-Tensor Imaging Findings and Cognitive Function Following Hospitalized Mixed-Mechanism Mild Traumatic Brain Injury: A Systematic Review and Meta-Analysis. Arch Phys Med Rehabil 2017; 11:2308–2319

30. Owen JP, Chang YS, Mukherjee P. Edge density imaging: mapping the anatomic embedding of the structural connectome within the white matter of the human brain. Neuroimage 2015; 109:402–17.

31. Owen JP, Marco EJ, Desai S, Fourie E, Harris J, Hill SS, Arnett AB, Mukherjee P. Abnormal white matter microstructure in children with sensory processing disorders. Neuroimage Clin 2013;2:844–53.

32. Owen JP, Wang MB, Mukherjee P. Periventricular White Matter Is a Nexus for Network Connectivity in the Human Brain. Brain Connect 2016;6:548–57.

33. Radhakrishnan R, Garakani A, Gross LS, Goin MK, Pine J, Slaby AE, Sumner CR, Baron DA. Neuropsychiatric aspects of concussion. Lancet Psychiatry 2016; (12):1166–1175.

34. Sato K, Kerever A, Kamagata K, Tsuruta K, Irie R, Tagawa K, Okazawa H, Arikawa-Hirasawa E, Nitta N, Aoki I, Aoki S. Understanding microstructure of the brain by comparison of neurite orientation dispersion and density imaging (NODDI) with transparent mouse brain. Acta Radiol Open 2017;6:2058460117703816.

35. Schneider T, Brownlee W, Zhang H, Ciccarelli O, Miller DH, Wheeler-Kingshott G. Sensitivity of multi-shell NODDI to multiple sclerosis white matter changes: a pilot study. Funct Neurol 2017;32(2):97–101.

36. Sepehrband F, Clark KA, Ullmann JF, Kurniawan ND, Leanage G, Reutens DC, Yang, Z. Brain tissue compartment density estimated using diffusion-weighted MRI yields tissue parameters consistent with histology. Hum Brain Mapp 2015;36:3687–702.

37. Smith SM. Fast robust automated brain extraction. Hum Brain Mapp. 2002; 17:143–55.

38. Timmers I, Roebroeck A, Bastiani M, Jansma B, Rubio-Gozalbo E, Zhang H. Assessing Microstructural Substrates of White Matter Abnormalities: A Comparative Study Using DTI and NODDI. PLoS One 2016;11:e0167884.

39. Wang MB, Owen JP, Mukherjee P, Raj A. Brain network eigenmodes provide a robust and compact representation of the structural connectome in health and disease. PLoS Comput Biol 2017;13:e1005550.

40. Xiao M, Ge H, Khundrakpam BS, Xu J, Bezgin G, Leng Y, Zhao L, Tang Y, Ge X, Jeon S, Xu W, Evans AC, Liu S. Attention Performance Measured by Attention Network Test Is Correlated with Global and Regional Efficiency of Structural Brain Networks. Front Behav Neurosci 2016;10:194.

41. Yang Y, Bender AR, Raz N. Age related differences in reaction time components and diffusion properties of normal-appearing white matter in healthy adults. Neuropsychologia 2015;66:246–58.

42. Yue JK, Vassar MJ, Lingsma HF, Cooper SR, Okonkwo DO, Valadka AB, Gordon WA, Maas AI, Mukherjee P, Yuh EL, Puccio AM, Schnyer DM, Manley GT; TRACK-TBI Investigators. Transforming research and clinical knowledge in traumatic brain injury pilot: multicenter implementation of the common data elements for traumatic brain injury. J Neurotrauma 2013;30:1831–44.

43. Yuh EL, Cooper SR, Mukherjee P, Yue JK, Lingsma HF, Gordon WA, Valadka AB, Okonkwo DO, Schnyer DM, Vassar MJ, Maas AI, Manley GT; TRACK-TBI INVESTIGATORS. Diffusion tensor imaging for outcome prediction in mild traumatic brain injury: a TRACK-TBI study. J Neurotrauma 2014;31:1457–77.

44. Yuh EL, Mukherjee P, Lingsma HF, Yue JK, Ferguson AR, Gordon WA, Valadka AB, Schnyer DM, Okonkwo DO, Maas AI, Manley GT; TRACK-TBI Investigators. Magnetic resonance imaging improves 3-month outcome prediction in mild traumatic brain injury. Ann Neurol 2013;73:224–35.

45. Zhang H, Hubbard PL, Parker GJ, Alexander DC. Axon diameter mapping in the presence of orientation dispersion with diffusion MRI. Neuroimage 2011;56:1301–15.

